# Cell-type specific analysis of physiological action of estrogen in mouse oviducts

**DOI:** 10.1101/2020.12.18.423483

**Authors:** Emily A. McGlade, Gerardo G. Herrera, Kalli K. Stephens, Sierra L. W. Olsen, Sarayut Winuthayanon, Joie Guner, Sylvia C. Hewitt, Kenneth S. Korach, Francesco J. DeMayo, John P. Lydon, Diana Monsivais, Wipawee Winuthayanon

## Abstract

One of the endogenous estrogens, 17β-estradiol (E_2_) is a female steroid hormone secreted from the ovary. It is well established that E_2_ causes biochemical and histological changes in the uterus. The oviduct response to E_2_ is virtually unknown in an *in vivo* environment. In this study, we assessed the effect of E_2_ on each oviductal cell type, using an ovariectomized-hormone-replacement mouse model, single cell RNA-sequencing (scRNA-seq), *in situ* hybridization, and cell-type-specific deletion in mice. We found that each cell type in the oviduct responded to E_2_ distinctively, especially ciliated and secretory epithelial cells. The treatment of exogenous E_2_ did not drastically alter the transcriptomic profile from that of endogenous E_2_ produced during estrus. Moreover, we have identified and validated genes of interest in our datasets that may be used as cell- and region-specific markers in the oviduct. Insulin-like growth factor 1 (*Igf1*) was characterized as an E_2_-target gene in the mouse oviduct and was also expressed in human Fallopian tubes. Deletion of *Igf1* in progesterone receptor (*Pgr*)-expressing cells resulted in female subfertility, partially due to an embryo developmental defect and embryo retention within the oviduct. In summary, we have shown that oviductal cell types are differentially regulated by E_2_ and support gene expression changes that are required for normal embryo development and transport in mouse models.

## Introduction

The oviduct (or Fallopian tube in humans) is a tube-like structure that connects the ovary to the uterus. The oviduct is divided into four main components known as the infundibulum, ampulla, isthmus, and the uterotubal junction (UTJ, also known as intramural junction in humans). The oviduct provides an optimal microenvironment for gamete fertilization, preimplantation embryo development, and embryo transport^1^. The majority of the cells encompassing the epithelial layer of the oviduct are non-ciliated (secretory) cells and ciliated cells. In the bovine oviduct, non-ciliated secretory cells are responsible for secreting a variety of proteins and metabolites to act as growth factors, and metabolic and immune regulators to aid in gamete maturation and fertilization^2^. Secretory cells are concentrated mainly in the proximal (isthmus and UTJ) region of the oviduct and are more sparsely dispersed toward the distal (infundibulum and ampulla) region of the oviduct^3^. However, ciliated cells are present in the opposing gradient compared to secretory cells: infundibulum>ampulla>isthmus ^3^. Ciliated cells are responsible for creating a flowing current of oviductal fluid toward the uterus that aids in proper embryo transport during early pregnancy^4^. Alteration of secretory function or inflammation of the oviduct, such as hydrosalpinx or salpingitis, negatively impacts the development of preimplantation embryos^5–7^. In addition, disruption of ciliary function has been shown to increase the risk for ectopic pregnancy (a condition where the embryo implants outside of the uterus)^8^. Therefore, the oviduct is crucial for the establishment of a successful pregnancy.

17β-estradiol (E_2_) is mainly produced from the granulosa cells of the ovary. In several mammalian species, elevated levels of E_2_ alter the histoarchitecture and function of the oviduct^9^. It is well-established that E_2_ is required for ovarian and uterine function. Global knockout of estrogen receptor α (*Esr1*^−/−^) in mice leads to female sterility, mainly due to anovulation and a uterine receptivity defect^10^. In other female reproductive tissues, such as the uterus, E_2_ induces the production of growth factors, e.g., insulin-like growth factor 1 (IGF1), in the stromal cell layer^11^. Subsequently, stromal-derived IGF1 then stimulates uterine epithelial cell production via a paracrine signaling pathway^12^. Interestingly, these growth factors (including IGF1, fibroblast growth factors (FGFs), complement component 3 (C3), and demilune cell parotid proteins (DCPPs)) have been identified as embryotrophic factors that facilitate preimplantation embryo development to the blastocyst stage and prevent apoptosis^13–17^. However, the mechanistic action and the fundamental understanding of E_2_ in the oviduct is virtually unknown. Our previous work showed that epithelial cell specific deletion of *Esr1* in the oviduct resulted in an alteration of protease and protease inhibitor production, leading to embryo death at 1.5 days post coitus (1.5 dpc)^18^. However, it is still unclear whether these E_2_-mediated proteases and protease inhibitors are expressed in specific epithelial cell types of the oviduct.

The use of single cell-RNA sequencing (scRNA-seq) analysis is an increasingly popular tool in the scientific community due to its capability to identify transcriptomes at a single cell resolution as well as to discover the heterogeneity of cell populations within tissues^19^. Single cell encapsulation for sequencing pipeline is commercially available^19^. In this study we used scRNA-seq analysis to determine the cell types and their response to E_2_ in the oviduct. Overall, the main objective of this study is to provide fundamental knowledge regarding the roles of E_2_ on the oviductal cellular response at a single cell resolution. Recent studies showed that the epithelial cells from the proximal and distal regions of the oviduct develop from two distinct cell lineages^20^. In contrast to these previous studies, our research focuses on the effect of E_2_ on all cell types present in the oviduct. Lastly, all genes expressed in our scRNA-seq datasets are readily available for the scientific community as a web search format.

## Materials and Methods

### Animals

All animals were maintained at Washington State University and were handled according to Animal Care and Use Committee guidelines using approved protocols #6147 and 6151. For scRNA-seq experiments, adult female mice (C57B6/J, 8-12 weeks old) were purchased from the Jackson (JAX) laboratory. Animals were ovariectomized and left for 14 days to recover and eliminate endogenous steroid hormones^12^. Sesame oil (100 μL) was used as a negative (Vehicle or Veh) control group. 17β-estradiol (E_2_) at a dose of 0.25 μg was diluted in 100 μL of 10% ethanol and sesame oil. Ovariectomized females were subcutaneously injected with Veh or E_2_. Twenty-four hrs after injection, the oviduct was collected for scRNA-seq analysis. Vaginal washes were collected from adult female mice (between 8-12 weeks old) to determine the stage of the estrous cycle^21^. At 11:00 h, oviducts from females in estrus were collected for subsequent scRNA-seq analysis.

The oviductal and uterine *Igf1* conditional knockout mouse model was generated by breeding progesterone receptor-Cre (*Pgr*^Cre/+^)^22^ with *Igf1*^f/f^ mice, purchased from Jax^23^. The conditional knockout females (*Pgr*^Cre/+^;*Igf1*^f/f^) and *Igf1*^f/f^ control littermates were used in the experiments. Genotyping protocols for *Pgr*^Cre^ and *Igf1*^f/f^ were performed as previously described^24^ and according to the recommendation from Jax for *Igf1*^f/f^. The cell-specific deletion of *Igf1* was validated and confirmed using *Igf1 in situ* hybridization (ISH) analysis using RNAscope, Cat# 443901: Mm-*Igf1* (Advanced Cell Diagnostics (ACD), Newark, CA) according to ACD recommended protocols.

### Histological, *in situ* hybridization (ISH), and immunohistochemistry (IHC) analysis

The oviduct and uterus were collected from female mice for histological analysis. Tissues were fixed in 10% formalin and processed for histological analysis as previously described^21^. Formalin-fixed oviductal and uterine tissues were paraffin embedded and sectioned to a 5-μm thickness. The antibodies used were anti-ESR1 antibody (Thermo Fisher Scientific, MA5-13191) at a dilution of 1:200, anti-PGR (Thermo Fisher Scientific, MA5-14505) at 1:400, and Ki67 antibody (BD Pharminogen, 550609) at 1:100. Vectastain ABC system (Vector Laboratories, Burlingame, CA) was used for colorimetric detection. ISH analysis was performed using RNAscope reagent kits and ACD HybEZ II Hybridization System. RNAscope probes #413281 Mm-*Dcn, #*583461 Mm-*Crabp2*, #583471 Mm-*Serpina1e*, and #583481 Mm-*Pdxk* were used for the detection of RNA in the oviductal tissues. Probes #313911 Mm-*Ppib* and #310043 *DapB* were used as positive and negative controls, respectively. Images were taken using a light microscope (Leica DMi8, Leica Microsystems, Buffalo Grove, IL).

### Quantification analysis of Ki67^+^ and PGR^+^ cells

Ki67^+^ and PGR^+^ cells were quantified using ImageJ software with Cell Counter Plugins. Ki67^+^ or PGR^+^ ciliated epithelial cells were counted and calculated as a percentage of total ciliated epithelial cells. A similar counting method was used for secretory and stromal cells (*n*=3–6 animals per treatment per group). The average total ciliated cell count for Ki67 analysis was 198 cells/animal. The range for ciliated cell count was 3-292 cells per microscopic field as there were very few ciliated cells present in the isthmus. The average total secretory and stromal cell counts for Ki67 analysis were 141 cells/animal (range: 98-185) and at 255 cells/animal (range: 120-418), respectively. The average total ciliated cell count for PGR analysis was 141 cells/animal (range:0-312 cells per microscopic field; again, as there was a very minimal number of ciliated cells present in the isthmus region). The average total secretory and stromal cell counts for PGR analysis were 160 cells/animal (range: 58-223) and at 271 cells/animal (range: 98-426), respectively.

### Oviductal cell sample preparation for scRNA-seq analysis

Oviducts were collected 24 hrs after Veh or E_2_ treatment (*n*=5-6 mice/group) or at estrus (*n*=5 mice). Oviducts were pooled and collected in Leibovitz-15 media (Gibco, 41300070, ThermoFisher Scientific, Carlsbad, CA) supplement with 1% fetal bovine serum (L15+1%FBS) and placed on ice. For Veh and E_2_-treated samples, the oviduct was dissected into distal (infundibulum with ampulla, referred to as InfAmp) or proximal (isthmus with UTJ, referred to as IsthUTJ) regions. To isolate oviductal cells, oviducts were quickly uncoiled by trimming the mesosalpinx in order to locate the ampulla-isthmus junction (AIJ). Then, the tissues were dissected at the AIJ region for subsequent InfAmp and IsthUTJ cell isolation. For estrus samples, the entire length of the oviduct was used. After dissection, 0.25% trypsin-EDTA (Sigma, T4049) was injected into the oviduct using a blunted 30-gauge needle until the trypsin was visibly flushed out from the other end of the oviduct. The tissues were then placed in a tube containing 500 μL of trypsin and incubated on ice for 30 min and at 37°C for another 30 min. After the incubation, 500 μL of FBS was added to stop the enzymatic activity of trypsin. Oviductal cells were then flushed out from oviductal tubes with 4°C L15+10%FBS. At this stage, the epithelial, stromal, and parts of muscle cell layers were visibly flushed out. Only translucent longitudinal muscle cells were left intact. Cell clumps and tissue debris were strained twice through 40-μm cell strainers. Cells were then spun down and resuspended in 0.04% bovine serum albumin AlbuMax (ThermoFisher Scientific, 11020-021) in phosphate-buffered saline. The final cell concentration was targeted for 8,000 cells/run.

scRNA-seq libraries were performed using the manufacturer’s protocol (10X Genomics Inc, Pleasanton, CA). Single Cell 3’ v2 chemistry was used. Briefly, individual cells (~8,000 cells/run) were separated into droplets by Gel bead in EMulsion (GEM) technology using 10X Chromium Controller. Emulsion beads were broken and barcoded cDNAs were pooled for amplification. Libraries generated were sequenced using an Illumina HiSeq4000, targeting 400M reads/run, paired-end, and 100bp read length.

### Human Fallopian tube collection

Human Fallopian tubes were collected at the Baylor College of Medicine with an Institutional Review Board approval number H-21138. Informed written consent was obtained from the patient prior to undergoing surgery. The tissue was collected from an individual without underlying gynecological diseases who underwent post-partum bilateral salpingectomy for sterilization. To obtain cell singlets, the Fallopian tubes were transected so that the luminal portion of the tube was exposed. The mucosal lining of the tubes was gently dissected with a scalpel and enzymatically digested in a solution of 0.05% Trypsin (Sigma, T1426) dissolved in HBSS (Gibco) for 30 minutes, followed by incubation in Collagenase (5 mg/ml, Sigma C5138) and DNAse I (200 μg/ml, Sigma DN25) solution for 20 minutes. The cells were filtered through a 40 μm filter, collected by centrifugation and resuspended in DMEM/F12 (Gibco, 11-330-057) supplemented with 10% FBS (Sigma, F2442) and antibiotics (Gibco, 15-240-096). Live cells were sorted on a BD FACS Aria II at the Cytometry and Cell Sorting Core Laboratory at Baylor College of Medicine (BCM) and submitted for single cell sequencing at the Single Cell Genomics Core Laboratory at BCM. scRNA-seq libraries were also performed using 10X Chromium Controller with Single Cell 3’ v3 chemistry. Libraries generated were sequenced using an Illumina NovaSeq 6000, targeting 300M reads/run, paired-end, 50bp read length.

### Processing of single cell RNA-seq data

The computer was equipped with AMD Ryzen 9 3900X 12-core, 24-thread and used as a server using Linux operating system. Processing of the raw sequencing data (mkfastq files) was carried out using Cell Ranger v3.1.0 to generate fastq files with default settings. Reference genomes, mm10-3.0.0 and GRCh38-3.0.0, were used for sequence alignment in mouse and human samples, respectively. Web summaries for each sample were shown in **Table 1**. Fastq files were then processed through Velocyto v0.1.24^25–27^ to produce loom files with spliced and unspliced mRNA information. Velocyto was run using the command run10x with default setting and a repeat masker gtf file from UCSC Genome Browser for mouse with an assembly for GRCm38/mm10 to mask expressed repeated elements. Generated loom files were then read in scanpy v1.6.0^28^ using JupyterLab^29^ for analyses and saved in an h5ad format^30^. For subsequent data analysis and visualization, a virtual machine (Docker)^31^ was used to create the analysis environment. Analysis packages including scanpy v1.6.0^28^, anndata v0.7.5^32^, umap v0.4.6^33^, numpy v1.19.4^34^, scipy v1.5.3^35^, pandas v1.1.4^36^, scikit-learn v0.23.2^37^, statsmodels v0.12.1^38^, python-igraph v0.8.3^39^, louvain v0.7.0^40^, and leidenalg v0.8.2^41^ were installed on the Docker.

**Table 1:**
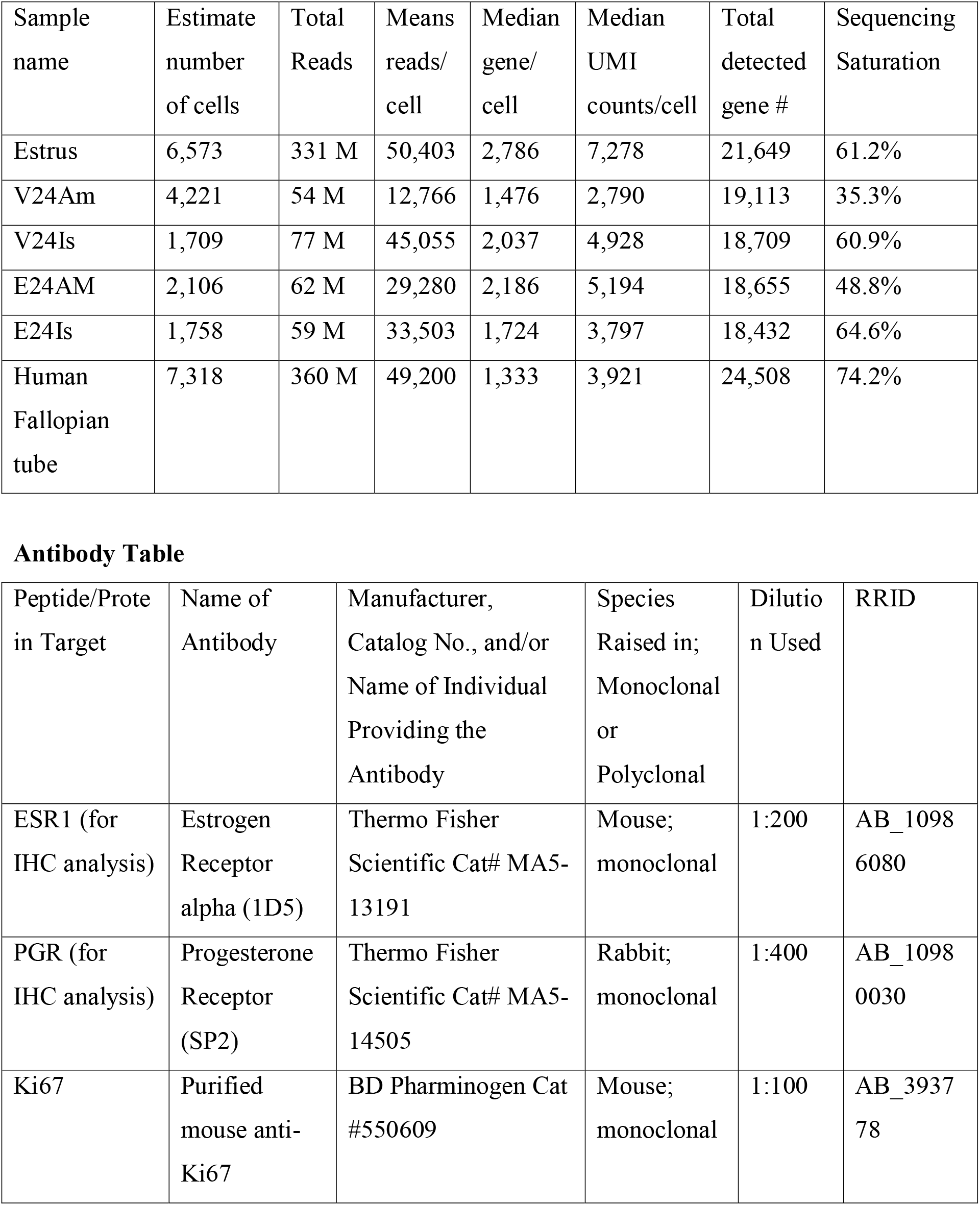
scRNA-seq output for each dataset in this study

| Sample name | Estimate number of cells | Total Reads | Means reads/cell | Median gene/cell | Median UMI counts/cell | Total detected gene # | Sequencing Saturation |
| --- | --- | --- | --- | --- | --- | --- | --- |
| Estrus | 6,573 | 331 M | 50,403 | 2,786 | 7,278 | 21,649 | 61.2% |
| V24Am | 4,221 | 54 M | 12,766 | 1,476 | 2,790 | 19,113 | 35.3% |
| V24Is | 1,709 | 77 M | 45,055 | 2,037 | 4,928 | 18,709 | 60.9% |
| E24AM | 2,106 | 62 M | 29,280 | 2,186 | 5,194 | 18,655 | 48.8% |
| E24Is | 1,758 | 59 M | 33,503 | 1,724 | 3,797 | 18,432 | 64.6% |
| Human Fallopian tube | 7,318 | 360 M | 49,200 | 1,333 | 3,921 | 24,508 | 74.2% |

**Antibody Table**
| Peptide/Protein Target | Name of Antibody | Manufacturer, Catalog No., and/or Name of Individual Providing the Antibody | Species Raised in; Monoclonal or Polyclonal | Dilution Used | RRID |
| --- | --- | --- | --- | --- | --- |
| ESR1 (for IHC analysis) | Estrogen Receptor alpha (1D5) | Thermo Fisher Scientific Cat# MA5-13191 | Mouse; monoclonal | 1:200 | AB_10986080 |
| PGR (for IHC analysis) | Progesterone Receptor (SP2) | Thermo Fisher Scientific Cat# MA5-14505 | Rabbit; monoclonal | 1:400 | AB_10980030 |
| Ki67 | Purified mouse anti-Ki67 | BD Pharminogen Cat #550609 | Mouse; monoclonal | 1:100 | AB_393778 |

### Quantifications and statistical analysis

Analysis was carried out using scanpy which was designed specifically for scRNA-seq data QC and analysis inspired by Seurat’s clustering tutorial^42^. Each sample was filtered out for doublets, genes that were present in less than 3 cells, cells that expressed less than 200 genes and cells that had higher than 5% reads mapped to mitochondrial genome. Samples were then processed through the default clustering vignette on scanpy including normalization, log1p, and scaling at default settings, ‘highly variable genes’ was set to 5000 genes, and Unique Molecular Identifier (UMI) and percentage mitochondrial were regressed out. Samples were then combined into one file using the concatenate function with batch keys for oviduct region when applicable. Principal component analysis (PCA), Uniform Manifold Approximation and Projection (UMAP) clustering analyses were then performed using 10 neighbors and PCs 1-37 based on variance ratio. Clustering was performed using Leiden clustering at 0.06-0.1 resolution based on known gene markers for secretory, ciliated, fibroblast, and muscle cells in the oviduct. Leiden clusters were then subsetted individually for independent analyses. Endothelial (*Pecam1^+^*) and immune cells (*Cd52^+^*) were filtered out from the dataset. When merging samples using the concatenate feature, there may be technical artifacts from sample collections making it difficult to interconnect different batches or samples. Batch balanced K-Nearest Neighbor (BBKNN)^43^ corrects these artifacts by using KNN to create connections between batches without interfering with PCA space or UMIs.

Using the generated Loom files from Velocyto, samples were processed following scVelo tutorials^27^; RNA Velocity Basics, Dynamic Modeling, and Differential Kinetics. The filter and normalize functions were set to 30 minimum shared counts and 10,000 top genes. The remaining functions were kept at default including moments. The RNA velocity map was then projected onto the UMAP and FA (ForceAtlas2)^44^ plots. FA plots are drawn from the function scanpy.tl.draw_graph() using default settings. Top 500 differentially expressed genes were used for determination of biological processes (BPs) enriched in each sample. These genes were uploaded onto geneontology.org using GO enrichment analysis with PANTHER overrepresentation test^45^ and Fisher test type with FDR correction when applicable.

### Data and code availability

All analyses in this study were saved in JupyterLab Notebook and deposited on github in https://github.com/winuthayanon/ovx_ve/ for the ovariectomized Veh- and E_2_-treated dataset, https://github.com/winuthayanon/estrus/ for the estrus dataset, and https://github.com/winuthayanon/humanFT/ for the human Fallopian tube dataset. Raw data as fastq files for the ovariectomized Veh- and E_2_-treated, the estrus, and the human Fallopian tube datasets were deposited at Gene Expression Omnibus (GEO accession numbers will be published upon acceptance of the manuscript).

### Embryo collections

Adult (8-12 weeks old) *Pgr*^Cre/+^;*Igf1*^f/f^ and *Igf1*^f/f^ female mice were bred overnight with C57B6/J male proven breeders. Females with a copulatory plug the next morning were considered pregnant at 0.5 days post coitus (dpc). Embryos were collected from the oviduct and uterus at 3.5 dpc (*n*=9-12 mice/genotype). Nonviable, unfertilized, and developmentally delayed embryos were also included in the total number of eggs/embryos. The region (oviduct vs. uterus) that the embryos were retrieved from was recorded. Images of embryos collected at 3.5 dpc were taken using Leica DMi8 microscope.

### Statistical analyses

All graphs represent mean◻±◻SEM. Individual value from each mouse was plotted when applicable. Statistical analysis was performed using GraphPad Prism v8.4.0 for Mac OS X (GraphPad Software, Inc., La Jolla, CA). Statistical significance is considered when *p*◻<◻0.05 using two-tailed unpaired Student’s *t*-test with Welch’s correction for simple comparison or two-way ANOVA with Sidak’s multiple comparisons test, unless otherwise indicated.

## Results

### E_2_-targeted gene expression in the oviduct

To determine the biological action of E_2_ on oviductal cell types, we ovariectomized adult female mice and treated with vehicle control (Veh) or E_2_ for 24 hrs. The concentration of 0.25 μg of E_2_ per mouse was chosen to allow a direct comparison between the oviductal response to E_2_ to another female reproductive tissue, the uterus. In our previous study, E_2_ concentration at 0.25 μg was shown to stimulate the proliferative response in uterine epithelial cells (**Fig. S1A**)^12^. We found that after treatment with E_2_, localization of ESR1 in the oviduct increased in the cytosolic compartment compared to the nucleus in the Veh-treated group (**Fig. 1A**), especially, in the epithelial cells of all regions including the infundibulum, ampulla, and isthmus. Images at a lower magnification are shown in **Fig. S1**. E_2_ treatment did not significantly change the number of Ki67^+^ ciliated or secretory cells (**Fig. 2B-C**). However, E_2_ significantly increased the number of Ki67^+^ stromal cells in the infundibulum. We also evaluated a known response of E_2_ treatment in the female reproductive tract, which causes an increase of progesterone receptor (PGR) expression in the stromal cell layer (**Fig. S1**)^12^. We found that E_2_ increased overall *Pgr* transcript levels in the whole oviduct (**Fig. 2D**). However, when we assessed protein levels using IHC analysis, we found that E_2_ exclusively induced PGR expression primarily in the ciliated cell population in the ampulla (**Fig. 1E-F**). As mentioned above, E_2_ induces the expression of IGF1 in the uterus^11^. Here we found that E_2_ increased *Igf1* transcript levels in the whole oviduct after 24 hrs of treatment (**Fig. 1G**). These data suggest that E_2_ treatment differentially induces cell specific proliferation and PGR expression in the oviduct but in a different fashion compared to the uterus (more detail in the discussion section).

**Figure 1.**
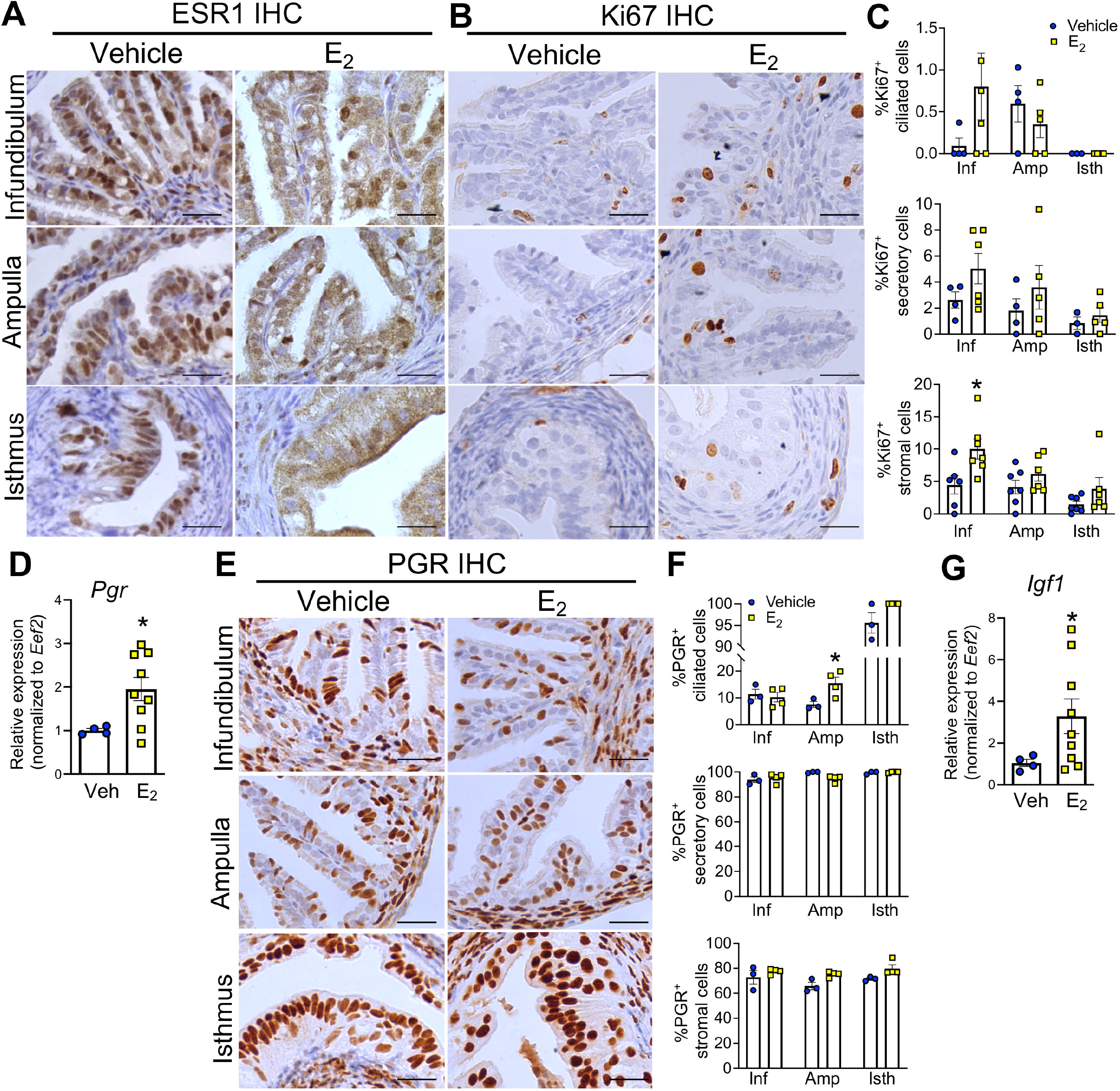
Ki67-positive cells and PGR expression in the mouse oviduct after E_2_ treatment. **A-B.** Immunohistochemical (IHC) analyses of ESR1 and Ki67 expression of the infundibulum, ampulla, and the isthmus of the oviduct, respectively. Oviducts were collected from ovariectomized mice after 24 hrs of treatment with Veh or E_2_ (0.25 μg/mouse), *n*=3-4 mice/group. **C.** Quantification of Ki67-positive ciliated, secretory and stromal cells. **D.** Transcript levels of *Pgr* in the whole oviduct after 24 hrs of Veh or E_2_ treatment (*n*=4-9 mice/group). **E.** IHC analysis of PGR expression after 24 hrs of treatment with Veh or E_2_. **F.** Quantification of PGR-positive ciliated, secretory and stromal cells. **G.** Transcript levels of *Igf1* in the whole oviduct after 24 hrs of Veh or E_2_ treatment (*n*=4-9 mice/group). Scale bars = 25 μm. Each dot represents data from each mouse. *, *p*<0.05, significantly different when compared to the corresponding Veh, unpaired Student’s *t*-test with Welch’s correction.

**Figure 2.**
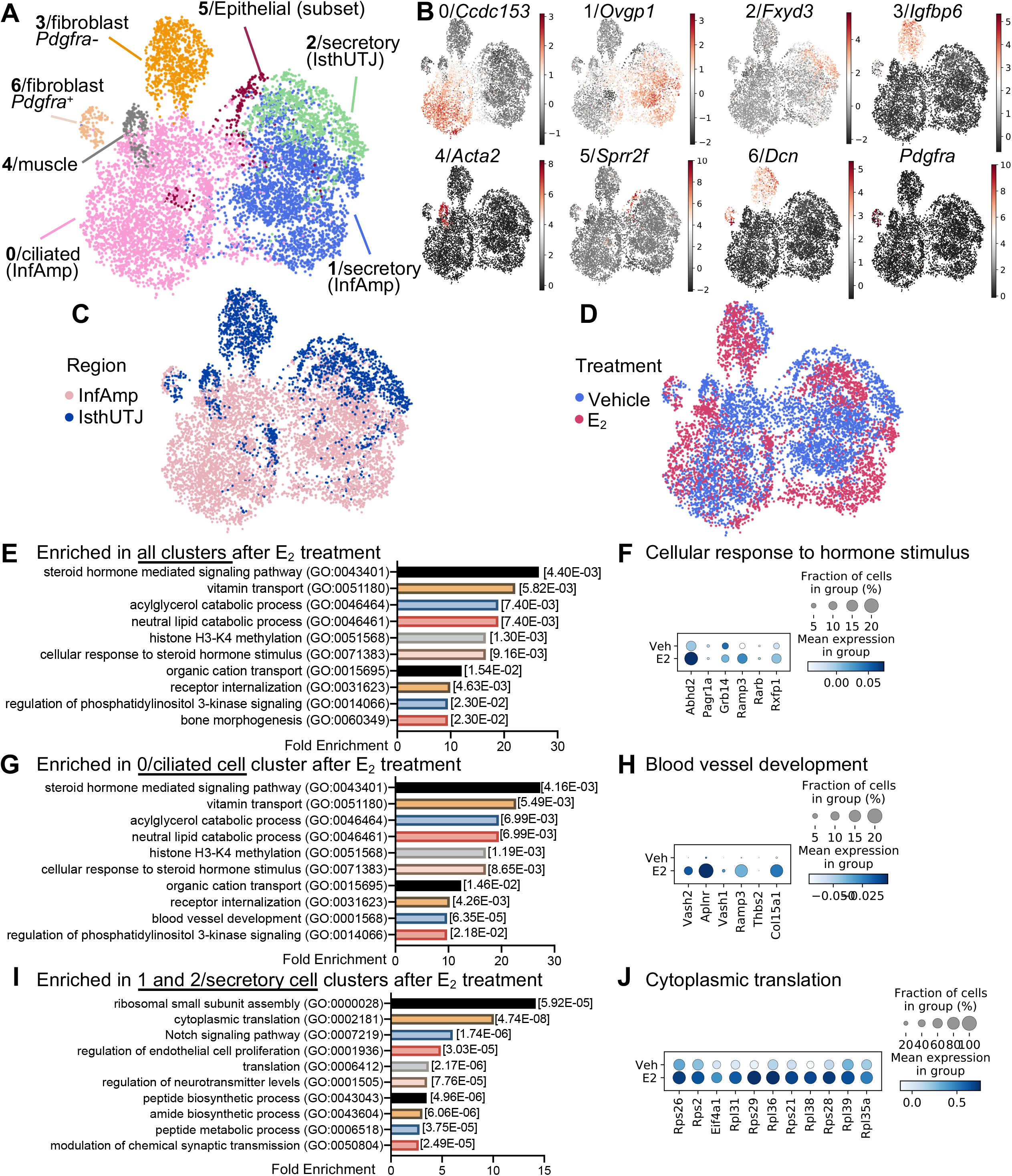
E_2_ treatment alters the transcriptional profile of the oviduct. **A.** Cell singlets were isolated from either the “infundibulum+ampulla” (InfAmp)” or “isthmus+uterotubal junction (IsthUTJ) and pooled from mice that were ovariectomized and treated with Veh or E_2_ for 24 hrs (*n*=5-6 mice/treatment). Single cell-RNA sequencing (scRNA-seq) data presented as Uniform Manifold Approximation and Projection (UMAP) plot. Each dot represents one cell. There are seven distinct oviductal cell clusters (#0-6) as indicated by different colors. **B.** Expression of the top marker gene from each cell cluster. **C-D**. All cell clusters differentiated by the region (InfAmp vs. IsthUTJ) or the treatment (Veh vs. E_2_). **E.** Top biological processes using gene ontology analysis from genes enriched in all clusters after E_2_ treatment. **F.** Dot plots of genes enriched in cellular response to steroid hormone stimulus in all clusters. Percentage of cells expressing particular gene(s) in the dataset is visualized by the size of the dot. Mean expression within each category is visualized by color. **G.** Top biological processes from genes enriched in 0/ciliated cell cluster after E_2_ treatment. **H.** Dot plots of genes enriched in blood vessel development in cluster 0/ciliated cells. **I.** Top biological processes from genes enriched in 1 and 2/secretory cell clusters after E_2_ treatment. **J.** Dot plots of genes enriched in cytoplasmic translation in clusters 1 and 2/secretory cells.

### E_2_-mediated transcriptional signatures in specific cell types in the oviduct

As shown above, E_2_ induced cellular responses in the oviduct in a cell-type- and region-specific manner. To delineate the gene regulatory effect of E_2_ in a cell-type- and region-specific manner, we employed scRNA-seq analysis. After 24 hrs of Veh or E_2_ treatment, oviducts were isolated into distal (infundibulum and ampulla, referred to as “InfAmp”) or proximal (isthmus and UTJ, referred to as “IsthUTJ”) regions and cell singlets were collected for scRNA-seq analysis. We found that cells from the oviduct were grouped into 7 cell clusters (**Fig. 2A**), including ciliated cells (InfAmp) indicated by pink and the expression of *Ccdc153* (**Fig. 2B**), secretory cells (InfAmp) indicated by purple and *Ovgp1*^+^, secretory cells (IsthUTJ) indicated by green and *Fxyd3*^+^, fibroblast (or stromal) *Pdgfra^-^* cells indicated by orange and *Igfbp6*^+^;*Dcn*^+^;*Pdgfra*^-^ (**Fig. S1A**), muscle cells indicated by grey and *Act2a*^+^, epithelial cells (subset) indicated by maroon and *Sprr2f*^+^, and fibroblast *Pdgfra*^+^ cells indicated by beige and *Dcn*^+^;*Pdgfra*^+^. The top 25 genes highly expressed in each cell cluster were shown in **Table S1**. In addition to cell clusters, we can also distinguish the regional origin (InfAmp vs. IsthUTJ) of each cell population (**Fig. 2C**). Our data showed for the first time that the transcriptional profile in each population was unique to the region. E_2_ treatment caused a shift of the transcriptome in all cell clusters (**Fig. 2D**). Specific genes in each cluster, region, or treatment in this dataset, can be searched on our laboratory website at www.winuthayanon.com/genes/ovx_ve/.

The effect of E_2_ and region-specific transcripts in each cell cluster are shown in dot plots (**Fig. S2B-M** and **Table S2** for top 500 genes). Using gene ontology (GO) analysis, we found that the top biological processes (BP) that were enriched in all cell clusters (as well as the ciliated cell clusters 0 and 6) after E_2_ treatment were steroid hormone mediated signaling pathways (GO:0043401, **Fig. 2E-G**). Examples of genes enriched in this cellular response to hormone stimulus included acylglycerol lipase (*Abhd2*), PAXIP1 associated glutamate rich protein 1 (*Pagr1a*), growth factor receptor-bound protein 14 (*Grb14*), receptor activity-modifying protein 3 (*Ramp3*), retinoic acid receptor beta (*Rarb*), and relaxin receptor 1 (*Rxfp1*) (**Fig. 2F**). One of the unique BPs in the ciliated cell clusters was blood vessel development, which included vasohibin 1 and 2 (*Vash1* and *Vash2*), apelin receptor (*Aplnr)*, *Ramp3*, thrombospondin 2 (*Thbs2*), and collagen type XV alpha 1 chain (*Col15a1*) (**Fig. 2H**). The top 3 BPs enriched in the secretory cell clusters included ribosomal small subunit assembly, cytoplasmic translation, and notch signaling pathway (**Fig. 2I**). We showed there that the BP ‘cytoplasmic translation’ enriched in E_2_-treated samples included ribosomal proteins S21, S26, S28, and S29 (*Rps21*, *Rps26*, *Rps28*, *Rps29*), ribosomal proteins L31, L35a, L36, L38, L39 (*Rpl31*, *Rpl35a*, *Rpl36*, *Rpl38*, *Rpl39*) and eukaryotic translation initiation factor 4A1 (*Eif4a1*). For a complete GOBP analysis for all cell clusters, refer to **Table S3**.

Based on our previous findings using microarray analysis, we found that proteases and protease inhibitors were regulated by E_2_- and ESR1-dependent action^18^. Here, we found that E_2_ treatment specifically increased overall expression of kallikrein 8 and 11 (*Klk8* and *Klk11*), prostatin (*Prss8*), neurotrypsin (*Prss12*), and serine protease 23 (*Prss23*) (**Fig. S3A**) in the oviduct. The expression of *Klk8* was specific to the secretory (IsthUTJ) cell cluster and *Klk11* to the ciliated (InfAmp) cell cluster (**Fig. S2B**). *Prss23* showed the highest expression among other proteases and the expression was specific to fibroblast *Pdgfra^-^* and muscle cell clusters. E_2_ increased overall expression of serine protease inhibitors (*Serpinb1a*, *Serping1*, and *Serpinh1*) and WAP four-disulfide core domain protein 18 (*Wfdc18* or extracellular peptidase inhibitor) (**Fig. S3C**). Similar to *Prss23*, protease inhibitors were expressed in fibroblast *Pdgfra^-^* and muscle cell clusters, but also in secretory and ciliated cells of the oviduct (**Fig. S3D**). These data suggest that E_2_ 1) modulates hormone signaling pathways in ciliated cells, 2) increases translation capacity in secretory cells, and 3) regulates the synthesis of proteases and protease inhibitors in multiple cell compartments of the oviduct.

### Insights into the effects of exogenous vs. endogenous E_2_ on oviductal cell responses

To determine whether exogenous E_2_ causes differential responses in oviductal cells compared to endogenous E_2_ at estrus, we compared the scRNA-seq data of oviductal cells collected at estrus to aforementioned E_2_-treated oviductal cells. Cells isolated during estrus were collected from the entire oviduct (denoted as “InfAmpIsthUTJ:estrus). scRNA-seq data were then combined and distinguished into InfAmp:E_2_-treated, IsthUTJ:E_2_-treated, and InfAmpIsthUTJ:estrus (**Fig. 3A**). We found that all 7 cell clusters present in the E_2_-treated group were also detected in the estrus samples (**Fig. 3B**). Colors for each cell cluster were consistent with the dataset from Veh- vs. E_2_-treated samples (**Fig. 2A**) for ease of comparison. Marker genes for each cell cluster are depicted in **Fig. S4**. Here, we show that endogenous E_2_ (estrus) had cell populations with similar transcriptome signatures compared to that of exogenous E_2_ (E_2_-treated) (**Fig. 3C**) as all cell populations were overlapped between estrus and E_2_-treated samples. Only one exception was that there were more secretory (IsthUTJ) cell populations in the estrus dataset compared to cells from the E_2_-treated samples.

**Figure 3.**
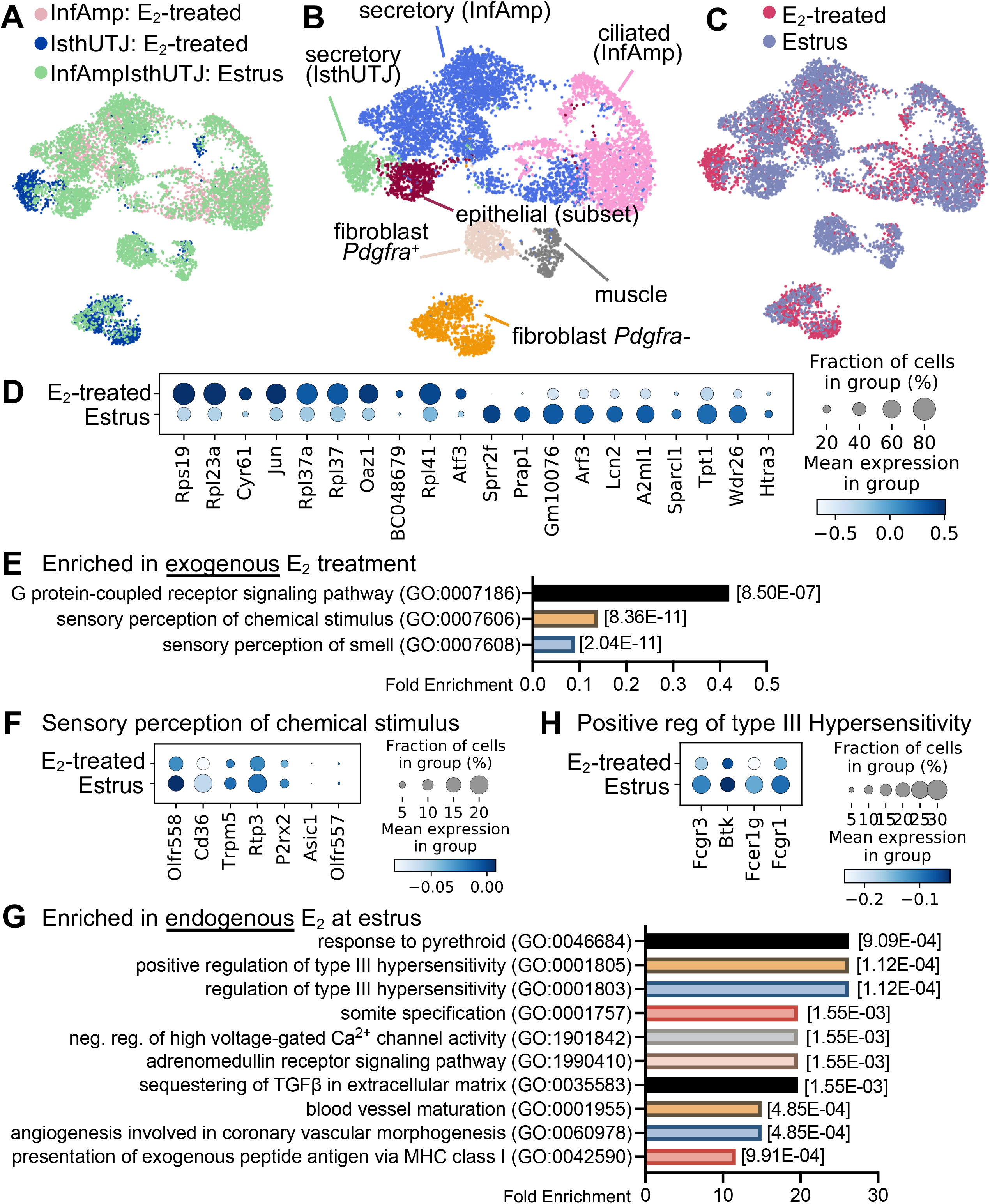
Exogenous and endogenos E_2_ induce differential transcriptional signatures in the oviduct. Endogenous E_2_ single cell samples were collected from the entire oviduct at the estrus stage (*n*=5 mice). scRNA-seq data from estrus (endogenous E_2_) were combined with data from ovariectomized E_2_-treated samples (exogenous E_2_) shown in Fig. 2 for comparison analyses. **A.** Estrus and E_2_-treated (InfAmp and IsthUTJ) datasets were overlapped. **B.** After the dataset from E_2_-treated and estrus samples were combined, scRNA-seq data were analyzed and separated into 7 cell clusters containing the similar cell populations to that in Fig. 2. **C.** Overall similarity and difference between endogenous vs. exogenous E_2_ in the oviductal cells. **D.** Dot plot of top 10 differentially expressed genes between exogenous and endogenous E_2_ in the oviduct. **E.** Top biological processes using gene ontology analysis of gene enrichment in exogenous E_2_ treatment. Dot plots of genes enriched in sensory perception of chemical stimulus in exogenous E_2_ treatment. **G.** Top biological processes of genes enriched in estrus samples. **H.** Dot plots of genes enriched in positive regulation of type III hypersensitivity in estrus samples.

We found that the top 10 genes elevated by E_2_ treatment were mostly ribosomal proteins including *Rpl19, Rpl23a, Rpl37, Rpl37a, Rpl41* (**Fig. 3D**), indicating a stimulation of transcription. Interestingly, common E_2_-target genes were also expressed at higher levels in the oviduct in E_2_-treated samples compared to estrus, such as cysteine rich angiogenic inducer 61 (*Cyr61*, aka IGF-binding protein 10)^46^ and *Jun*^47^. However, some other known E_2_-specific genes in the female reproductive tract were unique to estrus, such as small proline rich protein 2F (*Sprr2f*) and proline-rich acidic protein 1 (*Prap1*)^48,49^. Analysis using gene ontology demonstrated that E_2_-treated cells showed enrichment of molecules in G-protein-coupled receptor (GPCR) signaling pathways and both sensory perception of chemical stimulus and sensory perception of smell were the subset of GPCR signaling (**Fig. 3E-F**). Specific enriched genes in the sensory perception to chemical stimulus in E_2_-treated cells included olfactory receptor family 51 subfamily D and E member 1 (*Olfr557* and *Olfr558*), *Cd36*, transient receptor potential cation channel subfamily M member 5 (*Trpm5*), receptor transporter protein 3 (*Rtp3*), purinergic receptor P2X 2 (*P2rx2*), and acid sensing ion channel subunit 1 (*Asic1*). Endogenous E_2_ at estrus showed an enrichment of various BPs (**Fig. 3G**). As an example, transcripts enriched in the estrus sample for the BP of positive regulation of type III hypersensitivity (**Fig. 3H**) were Fc fragment of IgG receptor IIa, IgE receptor Ig, and IgG receptor Ia (*Fcgr3*, *Fcer1g,* and *Fcgr1*), and Bruton tyrosine kinase (*Btk*). Overall, these data indicate that the oviduct showed a slight difference in gene expression profile when exposed to E_2_ after ovariectomy compared to the estrus stage, at which time circulating E_2_ is peaked and progesterone is present at basal levels.

### Validation of markers identified from E_2_-treated InfAmp and E_2_-treated IsthUTJ samples

To validate the scRNA-seq findings observed from the ovariectomized E_2_-treated model, we used similar criteria to differentiate cell clusters in the scRNA-seq data from oviductal tissues collected at estrus. We found that all clusters found in E_2_-treated samples were present in the estrus sample (**Fig. 4A**). The top 10 marker genes for each cell cluster from the estrus dataset shown in the heatmap plot (**Fig. 4B**) were overlapped with the markers identified in Veh- and E_2_-treated samples (**Table S1**). Genes expressed in each cluster in this estrus dataset can be searched at www.winuthayanon.com/genes/estrus/.

**Figure 4.**
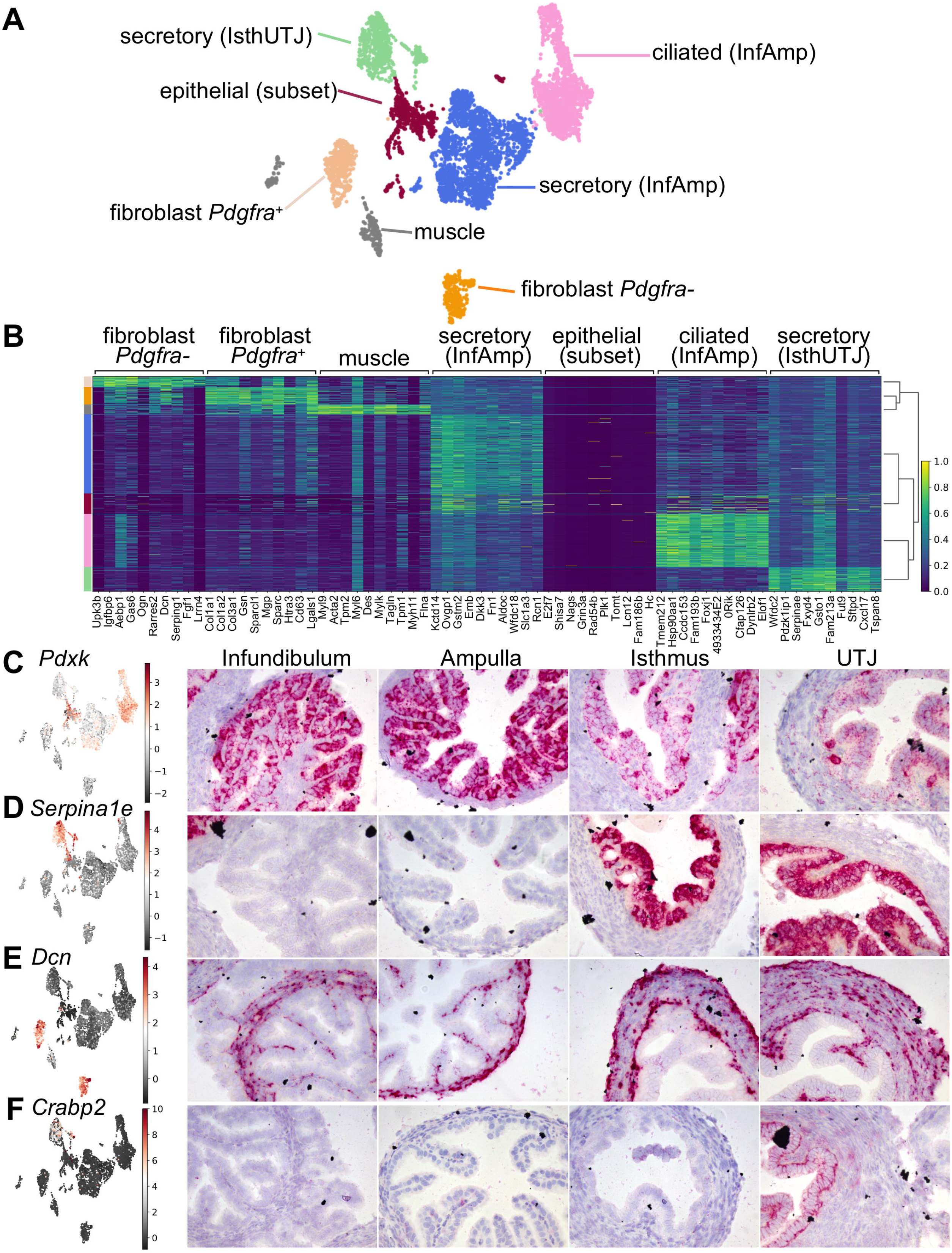
Cells isolated from estrus alone show similarity of cell clusters identified in the E_2_-treated dataset. **A.** UMAP plot of scRNA-seq analysis from estrus samples. Cells were separated into 7 cell clusters. **B.** Heatmap plot of top ten differentially expressed genes in each cell cluster. Color-coded bar on the left is coordinated with the cell clusters in A. *In situ* hybridization analyses of **C.** *Pdxk,* **D.** *Serpina1e*, **E.** *Dcn*, and **F.** *Crabp2* transcripts in the oviduct from samples collected randomly at different stages of the estrous cycle from ovarian intact adult female mice.

To verify the markers identified from each region from the E_2_-treated dataset (**Table S1**), we identified and determined the localization of these marker genes using ISH analysis in oviduct samples randomly collected during various stages of the estrous cycle. We found that pyridoxal kinase (*Pdxk*, also known as epididymis secretory sperm binding protein) was one of the top genes that was classified as an InfAmp-enriched gene (**Table S2**). Herein, we show that *Pdxk* was exclusively present only in the epithelial cells and detected at higher levels in the infundibulum and ampulla compared to isthmus and UTJ (**Fig. 4C**). Additionally, we were able to verify that *Serpina1e*, one of the top markers identified as IsthUTJ-specific, was only present in the epithelial cells in the isthmus and UTJ (**Fig. 4D**). Decorin (*Dcn*), a marker for fibroblasts (or stromal cells), was only detected in cells from mesenchymal origin including fibroblast (stroma) and muscle cell layers (**Fig. 4E**). Lastly, cellular retinoic acid binding protein 2 (*Crabp2*), was found specifically in the secretory (IsthUTJ) cluster and only detected in the epithelial cell layer of the UTJ region (**Fig. 4F**). Together, these data suggest that the markers identified from different populations and regions can be used as region- and cell-specific markers in the mouse oviduct.

### Embryotrophic factors, potential E_2_-target genes in the oviduct

One of the crucial functions of the oviduct is to support preimplantation embryo development. We showed that improper E_2_ signaling resulted in embryo death^18^. Embryotrophic factors were previously shown to improve embryo quality *in vitro*^13–17^. As *Igf1*, one of the embryotrophic factors identified, was upregulated by E_2_ (**Fig. 1G**), we looked more closely into the expression of embryotrophic factors in each cell population. We found that *Igf1* and its binding proteins (*Igfbp4*, *5*, and *6*) were mainly present in the cells from mesenchymal origin including fibroblasts (both *Pdgfra^-^* and *Pdgfra^+^*) and muscle cells (**Fig. 5A**). *Fgf1* and *Fgf9* were present in the fibroblast *Pdgfra^-^* population. However, *C3* was expressed in both fibroblast *Pdgfra^-^* and secretory (IsthUTJ) cell clusters. *Egf*, *Csf1*, and *Dcpp1-3* were expressed at lower levels and in a lower fraction of cells compared to *Igf1*, *Fgf1*, and *C3.* These findings indicate that embryotrophic factors are mainly present in cells originating from the mesenchyme. To determine if *IGF1*, *FGF1*, and *C3* were also expressed in human Fallopian tube, we isolated cells from one individual as a proof of concept. The heterogeneity of different cell populations within the Fallopian tube is shown in **Fig. 5B, S5**, and **Table S5**. *IGF1* was present in the fibroblast cell population (*DCN*+ and *PDGFRA*^+^). Interestingly, *FGF1* was detected only in a subset of fibroblasts. *C3* was also expressed in other cell types but at low levels. Specific genes in each cluster in this dataset, can be searched at www.winuthayanon.com/genes/humanFT/.

**Figure 5.**
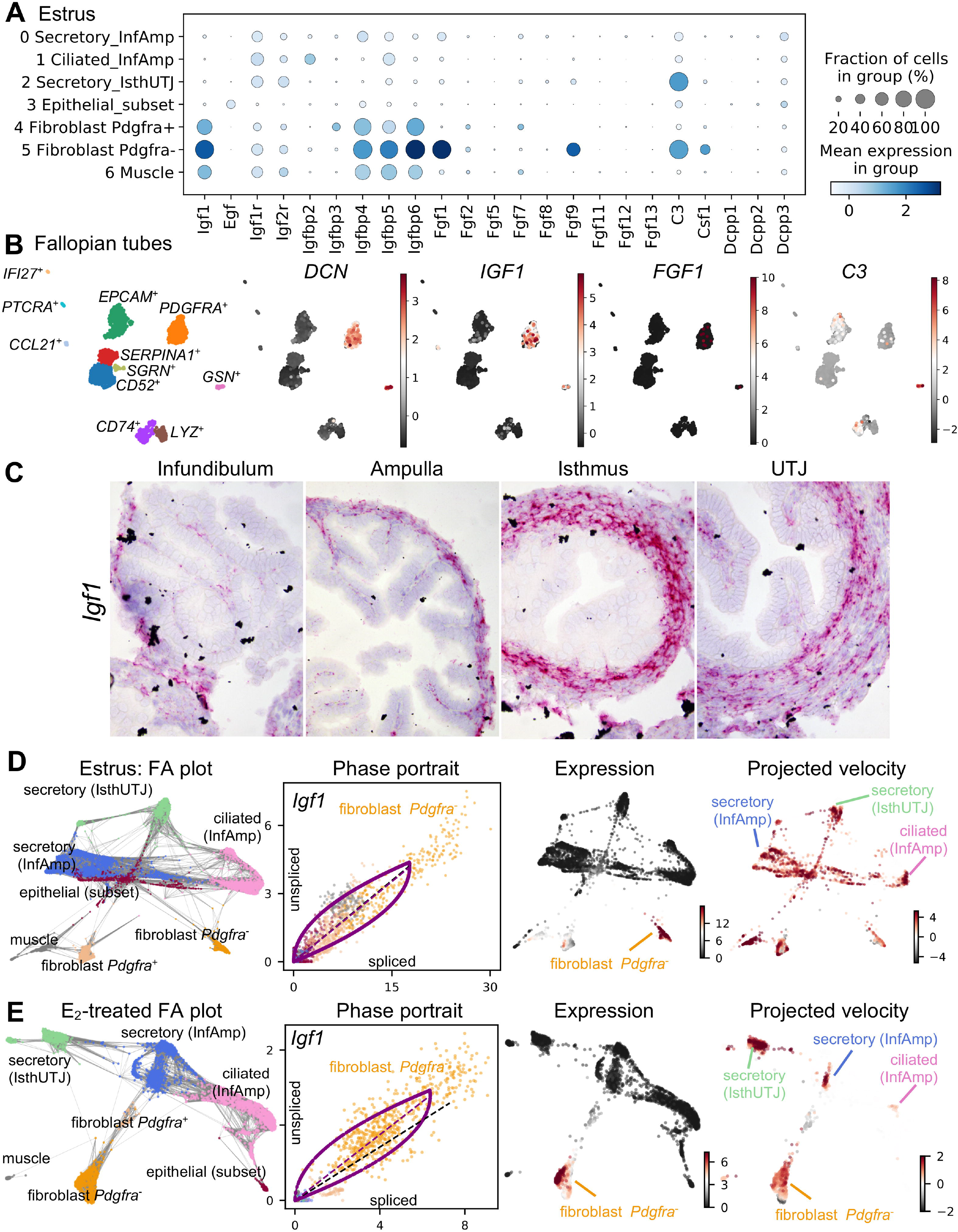
Unique cell populations in mouse oviducts and human Fallopian tube produce embryotrophic factor(s). **A.** Dot plot of embryotrophic factors and associated proteins in each cell cluster in the mouse oviduct collected at estrus. Insulin-like growth factor 1 (*Igf1*), epidermal growth factor (*Egf*), IGF receptor (*Igfr*), IGF-binding protein (*Igfbp*), fibroblast growth factor (*Fgf*), compliment C3 (C3), colony-stimulating factor 1 (*Csf1*), and demilune cell parotid proteins 1-3 *(Dcpp*). Different sizes of the circle represent percentages of the cells with expression of each gene. Color scale represents normalized expression value. **B.** scRNA-seq analysis of cell clusters from human Fallopian tube isolated from one individual as a proof-of-concept. Marker genes for each cell cluster is indicated in Figure S5. Stromal cells (decorin, *DCN*+) also express *IGF1, FGF1*, and *C3.* **C.** Expression of *Igf1* in the mouse oviduct using ISH analysis. **D.** and **E.** ForceAtlas2 (FA) plot, phase contrast, expression, and RNA project velocities of *Igf1* from mouse oviductal cells at (**D**) estrus or (**E**) after E_2_ treatment. Gray lines in the FA plot represent all velocity-inferred cell-to-cell connections/transitions but does not differentiate the directionality of velocity. Phase portraits depict a spliced vs. unspliced *Igf1* mRNA. Black dotted line represents constant transcriptional state. The expression of *Igf1* is mainly in the fibroblast cell populations, but the inferred direction of RNA velocity (projected velocity) showed the trajectory towards ciliated (InfAmp), secretory (InfAmp), and secretory (IsthUTJ) cell populations.

As IGF1 was present at high levels in both mouse oviducts and human Fallopian tubes, we reasoned that IGF1 was a viable candidate as an E_2_-target gene in the oviduct and could provide biological potential for oviductal function. First, we validated the expression of *Igf1* in the mouse oviduct using ISH analysis, as there was no reliable IGF1 antibody. Here, we showed that *Igf1* was indeed expressed in the fibroblasts and muscle cell layers of the oviduct (**Fig. 5C**) and the expression in the epithelial cells was omissible.

To determine gene regulation trajectories, we used RNA velocity analysis to assess the spliced vs. unspliced *Igf1* mRNA. RNA velocity of particular genes can be used to predict the future state (inferred direction) of individual cells based on the time derivative of splicing information and gene expression dynamic^25,50^. We identified *Igf1* as one of the driving genes of the transcriptional dynamic in both estrus and E_2_-treated datasets (**Fig. 5D-E**). Grey lines in the FA plots represent all velocity-inferred cell-to-cell connections but does not differentiate the directionality of velocity. Phase portraits showed the ratio of spliced vs. unspliced *Igf1* mRNA in each cell cluster (color-coded dots) in estrus (**Fig. 5D**) and E_2_-treated (**Fig. 5E**) samples. *Igf1* was strictly expressed in the fibroblast *Pdgfra^-^* population but had an inferred direction (projected velocity) upon epithelial cell clusters such as secretory (InfAmp), secretory (IsthUTJ), and ciliated (InfAmp). These data indicate that IGF1 is expressed in the fibroblast population in both mouse and human tissues and *Igf1* has a potential impact on gene regulation in other cell types within the mouse oviduct.

### *Loss of* Igf1 *expression led to a partial embryo developmental defect and embryo retention in the oviduct*

To test whether IGF1 is functionally required for female reproduction, we conditionally ablated *Igf1* expression in *Pgr* expressing cells using the *Pgr*^Cre/+^*;Igf1*^f/f^ mouse model. We validated that *Igf1* transcript levels were significantly reduced in *Pgr*^Cre/+^*;Igf1*^f/f^ compared to *Igf1*^f/f^ control oviduct (**Fig. 6A**). The overall histoarchitecture of the oviduct appeared to be similar between *Pgr*^Cre/+^*;Igf1*^f/f^ and of *Igf1*^f/f^ mice (**Fig. 6A**). Previous studies showed that *Pgr*^Cre/+^*;Igf1*^f/f^ females were subfertile^51^. However, it is still unclear if the fertility defect was due to a direct disruption in oviductal function. In our study, we showed that morula and blastocyst stage embryos were collected from *Igf1*^f/f^ females at 3.5 days post coitus (dpc) (**Fig. 6B-D**). We found that developmentally delayed or nonviable embryos were present in the *Pgr*^Cre/+^*;Igf1*^f/f^ females, but not significantly different than those collected from *Igf1*^f/f^ mice (**Fig. 6D**, *p=*0.054). The number of total eggs or embryos was similar between *Pgr*^Cre/+^*;Igf1*^f/f^ and *Igf1*^f/f^ females. Nevertheless, there were significantly more embryos retained in *Pgr*^Cre/+^*;Igf1*^f/f^ oviduct compared to *Igf1*^f/f^ females (**Fig. 6F**). Moreover, these embryos retained in the oviduct appeared to be developmentally delayed, unfertilized, or nonviable (**Fig. 6G**). These findings suggest that a loss of IGF1 could lead to an embryo developmental defect and embryo retention in the oviduct.

**Figure 6.**
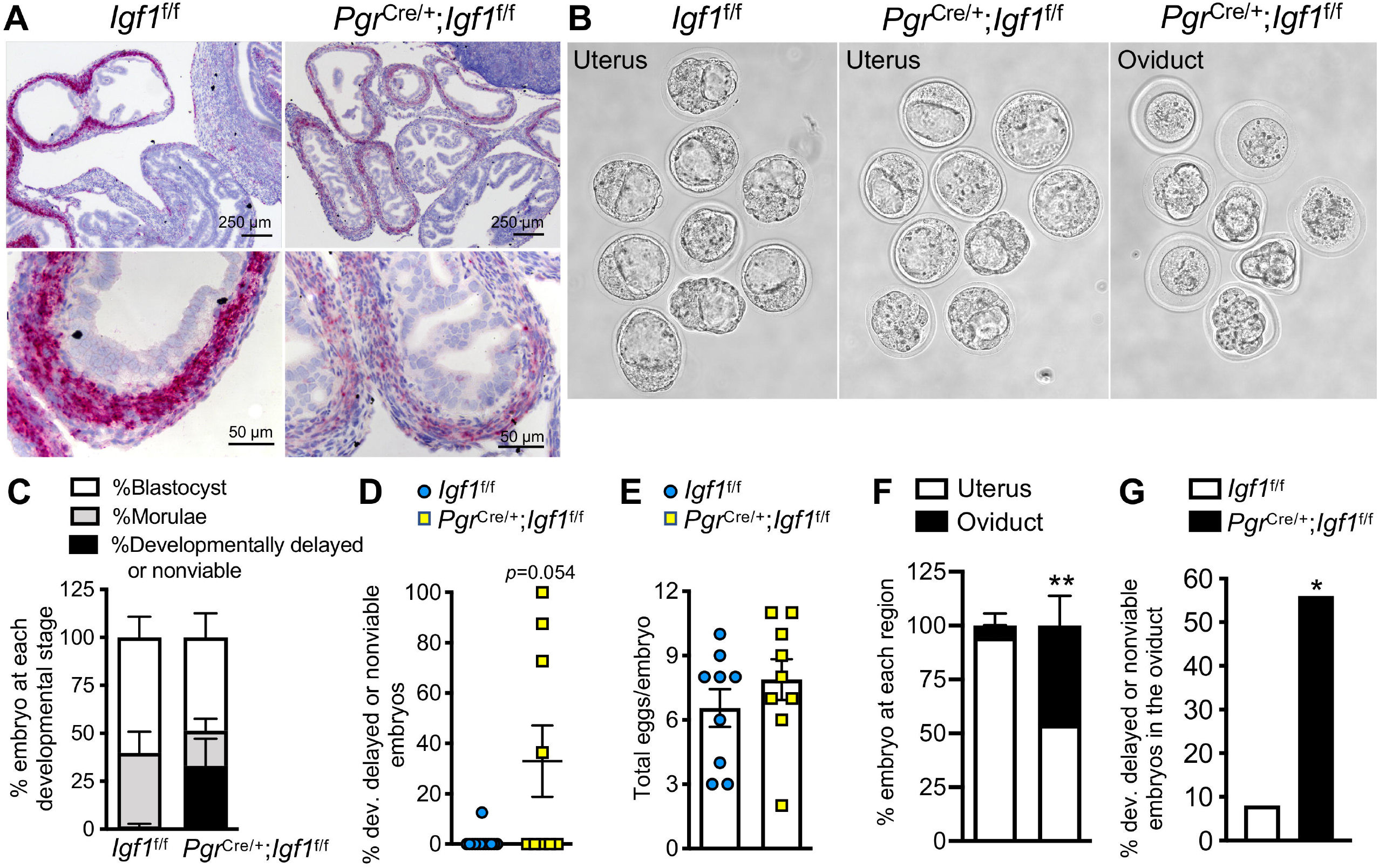
Oviductal ablation of *Igf1* increases developmentally delayed and nonviable embryos in mouse oviducts and inhibits embryo transport. **A.** *Igf1* transcripts in *Igf1*^f/f^ (control) compared to *Pgr*^Cre/+^;*Igf1*^f/f^ oviducts. **B.** Embryos collected from the uterus and/or oviduct from *Igf1*^f/f^ compared to *Pgr*^Cre/+^;*Igf1*^f/f^ females at 3.5 days post coitus (dpc). **C.** Percentage of embryos at each developmental stage at 3.5 dpc. **D**. Percentage of developmentally delayed or nonviable embryos. *p=*0.054, unpaired Student’s *t-*test with Welch’s correction. **E**. Total number of all eggs or embryos present in the oviduct at 3.5 dpc. **F.** Percentage of embryos collected at each region of the oviduct at 3.5 dpc. **, *p*<0.01, significantly different when compared to the corresponding *Igf1*^f/f^ mice, unpaired Student’s *t*-test with Welch’s correction. **G.** Percentage of mice with developmentally delayed or nonviable embryos in the oviduct. *, *p*<0.05, significantly different when compared to the corresponding *Igf1*^f/f^ group, Fisher’s exact test for contingency plot. *n=*9-12 mice/genotype.

## Discussion

In the uterus, E_2_ recruits all epithelial cells to undergo proliferation^46^. However, in the oviduct E_2_ showed minimal effect on epithelial cell proliferation. Moreover, the expression pattern of PGR in the oviduct was distinct from that of the uterus. In the uterus, PGR is mainly present in the epithelial cells in the absence of E_2_. Upon E_2_ treatment, uterine expression of PGR is decreased in the epithelial cells but increased in the stromal and muscle cell layers^52^. Although developing from the same fetal origin, the Müllerian duct, the oviduct responds to steroid hormones in a distinct manner compared to the uterus. We found that PGR was detected in almost 100% of secretory cells regardless of the oviductal region. The expression of PGR in secretory cells and stromal cells was not altered in the presence or absence of E_2_. However, E_2_ significantly increased PGR^+^ ciliated cells in the ampulla, suggesting that E_2_-induced PGR expression in the oviduct is exclusively specific to ciliated cells. These data indicate that distinct epithelial cell populations, e.g., ciliated and secretory cells, are differentially regulated by steroid hormone signaling, potentially in a region-specific manner.

Using scRNA-seq analysis, we found that transcriptional signatures of both epithelial cells and fibroblast were altered after E_2_ treatment. Interestingly, the ciliated cell cluster’s top 1 BP was ‘steroid hormone mediated signaling pathway’ whereas the secretory cell (from both InfAmp and IsthUTJ) clusters’ top two BPs were ‘ribosomal small subunit assembly’ and ‘cytoplasmic translation’. This further indicates that ciliated and secretory cells of the oviduct respond differentially to E_2_ in a distinguishable cell-type specific manner. After E_2_ treatment, secretory cells were enriched for genes responsible for translation. Not surprisingly, exogenous E_2_ caused a slight shift in the transcriptome of oviductal cell types from that of endogenous E_2_ during estrus. These differences are likely due to the presence of progesterone at a basal level at the estrus stage in contrast to the absence of progesterone when the mice were ovariectomized and treated with exogenous E_2_. However, the majority of the transcriptome signatures remained similar as indicated by overlapping cells when comparing estrus and E_2_-treated samples.

In addition to previous studies^20^, our scRNA-seq analysis from Veh- and E_2_-treated samples revealed that there are other genes that can be used as region- and cell-type-specific markers. We found that *Wt1* was expressed in the infundibulum and ampulla (**Table S2**), consistent with previous reports^20^. Here, we also showed that *Pdxk* was highly expressed in all epithelial cell types in the infundibulum and ampulla. *Pdxk*, is also expressed in the bovine oviduct^53^, suggesting that its expression in other mammalian species might be region-specific as well. *Serpina1e* was detected exclusively in the secretory cells of the oviduct in the isthmus and UTJ. *Dcn*, a small leucine-rich pericellular matrix proteoglycan, was expressed in cells from mesenchymal origin including fibroblast and muscle cells in both mouse oviducts and human Fallopian tubes. We also identified that *Crabp2* was expressed in epithelial cells of the UTJ. Mice with biallelic knockout of five *Serpina1a–e* develop emphysema but showed no reproductive phenotype^54^. *Dcn*^−/−^ and *Crabp2*^−/−^ mice are also viable and fertile^55,56^. These findings suggest that a loss of these genes do not affect female fertility. Therefore, *Serpina1e*, *Dcn*, and *Crabp2* can be specifically targeted for Cre allele knock-ins as a tool for cell-type- and region-specific deletion of genes of interest in the oviduct.

Evidence showed that supplementation of culture media with embryotrophic factors increases embryo quality and the number of blastocysts produced *in vitro*^13-17,57^. Previous studies showed that deletion of *Igf1* affects female fertility in mice^51^. The reproductive defect in *Pgr*^Cre/+;^*Igf1*^f/f^ mice did not seem to emanate from a uterine phenotype^51^. However, it was still unclear whether the *Pgr*^Cre/+;^*Igf1*^f/f^ mice had an oviductal defect. Here, we illustrated that a loss of *Igf1* in the oviduct resulted in a trending increase in the number of embryos with delayed development or nonviable embryos. Although the number was not significantly different from the control group, it is astounding that a lack of a single factor, *Igf1*, could result in a detectable defect in preimplantation embryo development. One possibility could be that liver-derived circulatory IGF1 present in the oviductal fluid could also act as an additional source of IGF1 for the embryos in the *Pgr*^Cre/+;^*Igf1*^f/f^ mice. Nevertheless, our findings provide evidence that the oviduct provides a finely tuned balance for embryos to develop normally in the oviduct. In addition to an embryo developmental defect, we also found that these developmentally delayed, nonviable embryos, or unfertilized eggs were retained in the oviduct *in vivo*. This phenomenon has also been observed in mares^58–60^, hamsters^61^, and rats^62^.

As the IGF1 receptor is also present on bovine embryos^63,64^, it is possible that IGF1 secreted from the oviduct promotes a positive-feedback loop to the embryo in a paracrine fashion in mammals. In the IVF setting, human embryos were often cultured in the media supplemented with growth factors^65^. Therefore, this paracrine signaling of IGF1 likely supports preimplantation embryo development in mammalian species, including humans. Although the presence of developmentally delayed or nonviable embryos in the *Pgr*^Cre/+;^*Igf1*^f/f^ was not significantly different from *Igf1*^f/f^ mice, we found that IGF1 provided a non-negligible function in the oviduct for proper embryo development. Future studies could address whether C3 is required for normal embryo development as C3 was another embryotrophic factor identified in both mouse oviducts and human Fallopian tubes. In addition, more human samples are required to characterize the transcriptome and heterogeneity of cell populations within the Fallopian tube.

In summary, we showed the heterogeneity of epithelial cell and fibroblast populations in the oviduct. Each cell type distinctly responded to the presence of the steroid hormone, E_2_. Not surprisingly, oviductal cells behaved differently from that of the uterus (or the mammary gland^66^) in response to E_2_, suggesting a tissue-specific regulation of steroid hormones within the female reproductive tract in mammals. The use of scRNA-seq analysis revealed that supraphysiological treatment of E_2_ triggers a relatively similar response in oviductal cell populations to that of endogenous E_2_ during the estrus stage. We found that IGF1 is a potential candidate as an E_2_-target gene in the oviduct and is necessary for normal embryo development and transport. Therefore, IGF1 could potentially be an endpoint marker for diagnosis of problematic pregnancies in humans. Our findings suggest that E_2_ regulation of oviductal cell response and function could provide crucial support for preimplantation embryo development and transport in mammals.

## Acknowledgement

The authors thank Yu Ishikawa for oviductal cell isolation protocol; Anna Willie, Jeffery Erickson and Jalene Velezques for some of the immunohistochemical staining; and Ryan Driskell for initial assistance with scRNA-seq data analysis pipeline.

## Author Contributions

E. McGlade, G. Herrera, and W. Winuthayanon designed research; E. McGlade, G. Herrera, K. Stephens, and S. Olsen, analyzed data; E. McGlade, G. Herrera., K. Stephens, S. Olsen, J. Guner, S. Hewitt, D. Monsivais, and W. Winuthayanon performed research; E. McGlade, G. Herrera, and W. Winuthayanon wrote the paper; F. DeMayo, J. Lydon, and K. Korach contributed new reagents or analytic tools; and G. Herrera and S. Winuthayanon developed software necessary to perform and record experiments.

