## Supplementary materials for "Cell-type specific analysis of physiological action of estrogen in mouse oviducts"

**Supplementary figures:**

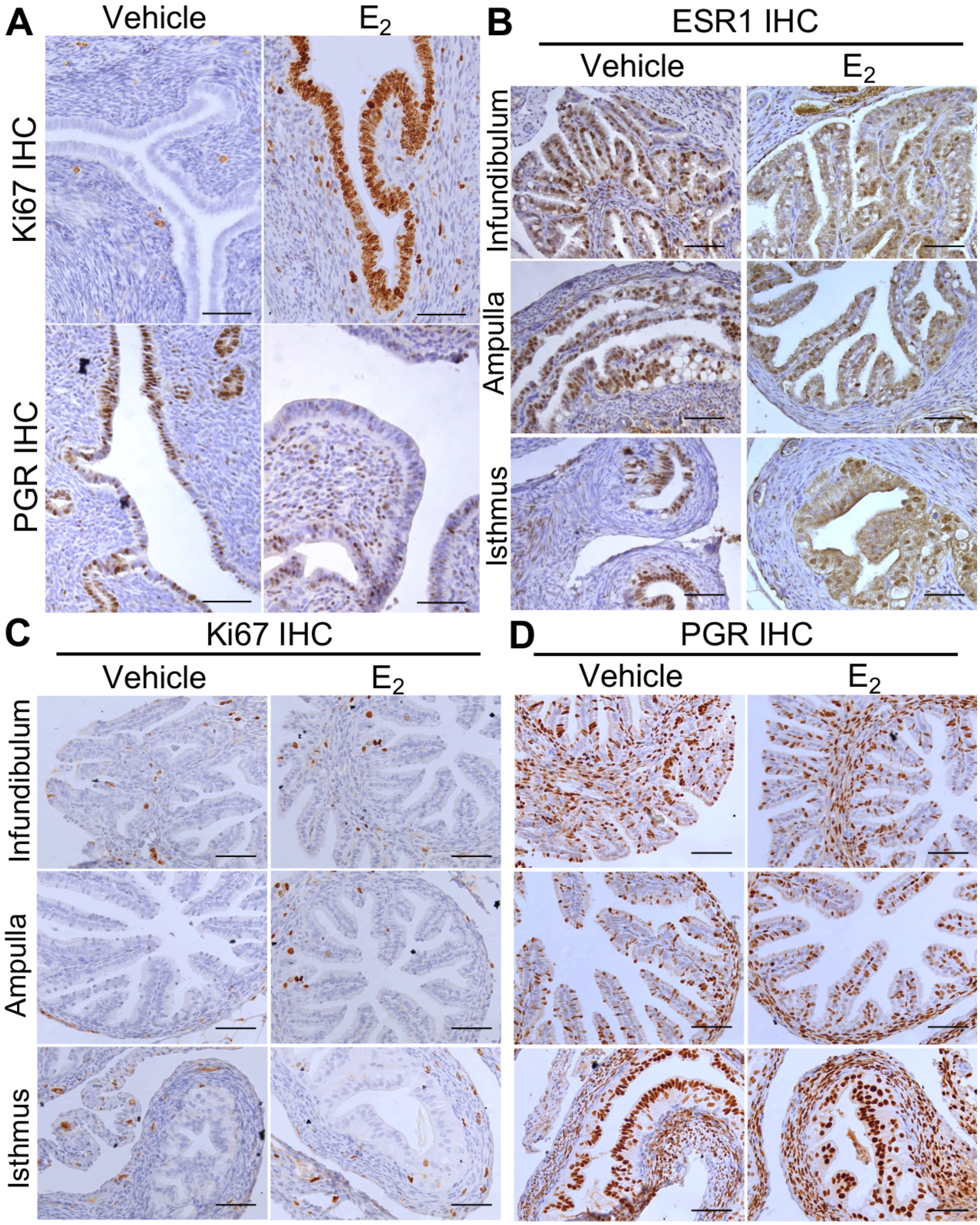


**Figure S1.** IHC Staining of ESR1, Ki67, and PGR in uterine and oviduct samples collected from ovariectomized females treated with Veh or $E_2$. **A.** Staining of Ki67 and PGR in uterine samples after Veh or $E_2$ treatment. **B-D.** Lower magnification of ESR1, Ki67, and PGR staining images from Fig. 1 of oviduct samples treated with Veh or $E_2$. All scale bars = 50 μm.

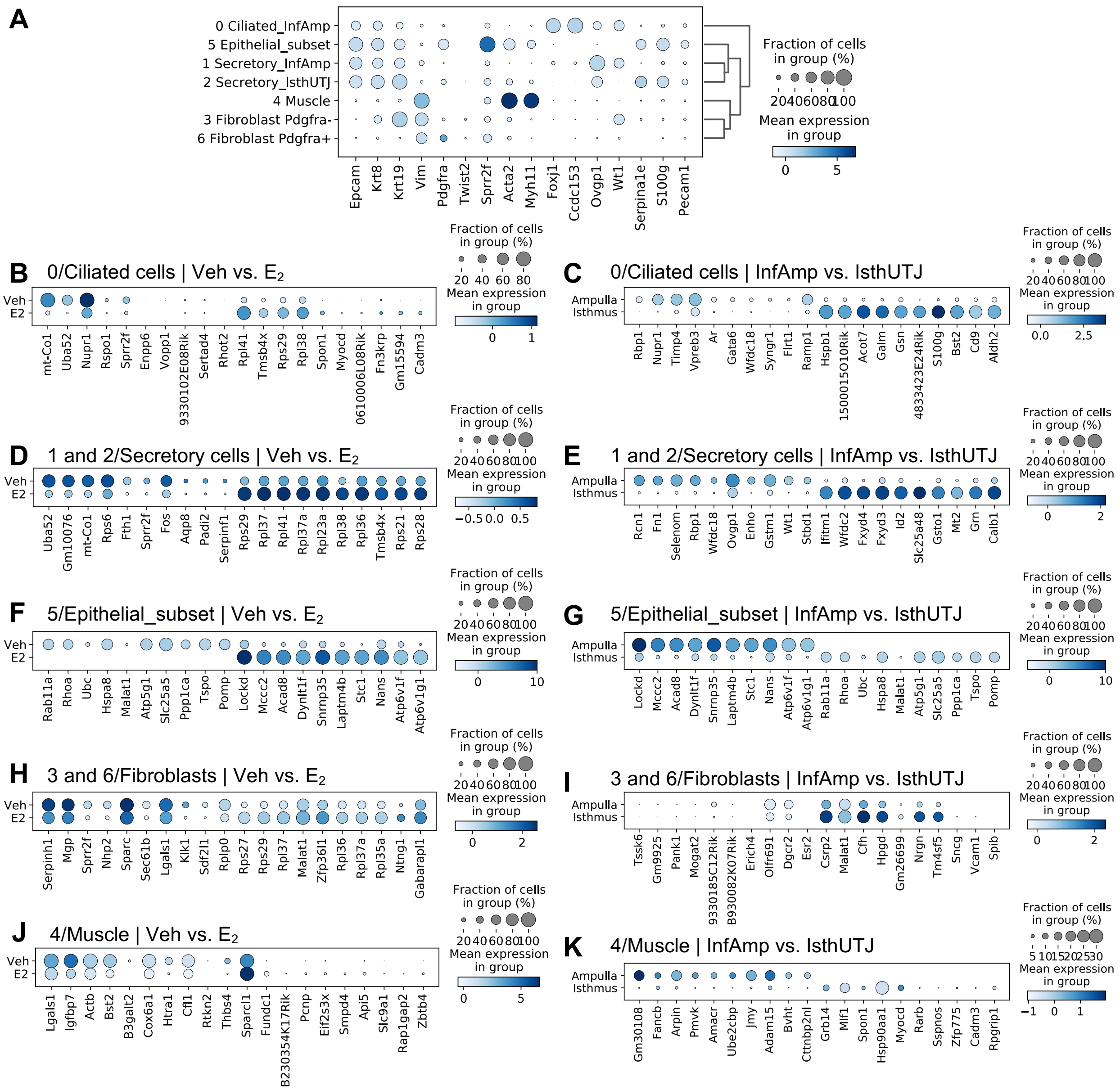


**Figure S2.** Dot plots of top 10 differentially expressed genes in cell clusters comparing between Veh vs. $E_2$ treatment or InfAmp vs. IsthUTJ. **A.** Marker genes in all cell clusters including 0/ciliated (InfAmp), 1/secretory (InfAmp), 2/secretory (IsthUTJ), 3/fibroblast *Pdgfra*$^-$, 4/muscle, 5/epithelial (subset), and 6/fibroblast *Pdgfra*$^+$ cell clusters. Epithelial cells are indicated by *Epcam*$^+$ and *Krt8*$^+$, mesenchymal cells (fibroblasts and muscles) by *Vim*$^+$, *Pdgfra*$^+$, and *Twist2*$^+$, muscle cells by *Act2a*$^+$ and *Myh11*$^+$, ciliated cells by *Foxj1*$^+$ and *Ccdc153*$^+$, secretory cells by *Ovgp1*$^+$, and endothelial cells by *Pecam1*$^+$. *Krt19* was expressed in both epithelial and fibroblasts. *Sprr2f*, *Serpina1e*, and *S100g* were expressed specifically in 5/epithelial cell subset. **B-K**. The top 10 region- and $E_2$-specific genes for each cell clusters are shown.

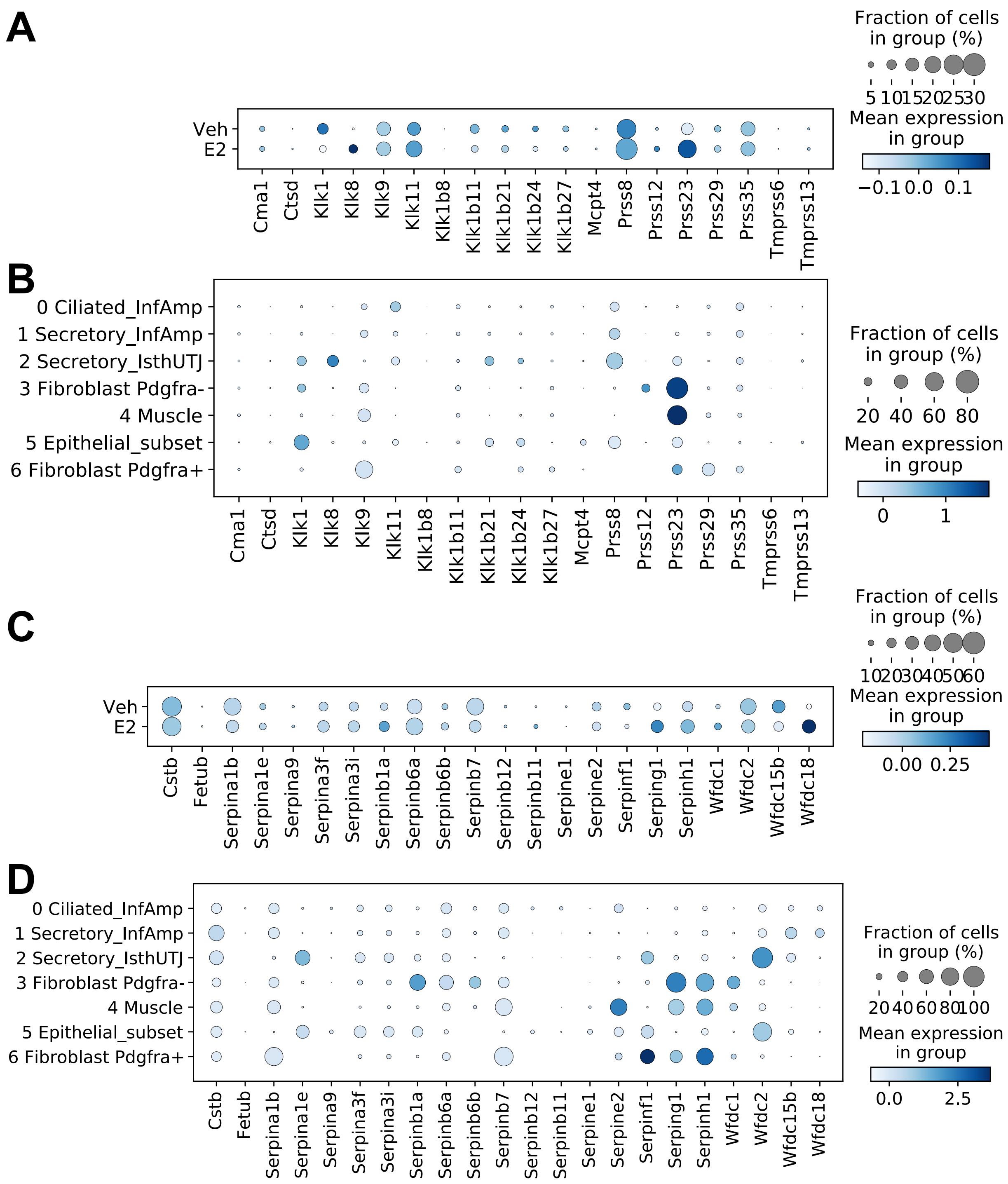


**Figure S3:** Dot plots of genes in (**A-B**) protease and (**C-D**) protease inhibitor family in cells isolated from Veh and E2-treated samples; including 0/ciliated, 1/secretory (InfAmp), 2/secretory (IsthUTJ), and 3/fibroblast *Pdgfra*-, 4/muscle, 5/epithelial (subset), and 6/fibroblast *Pdgfra*+ cell clusters.

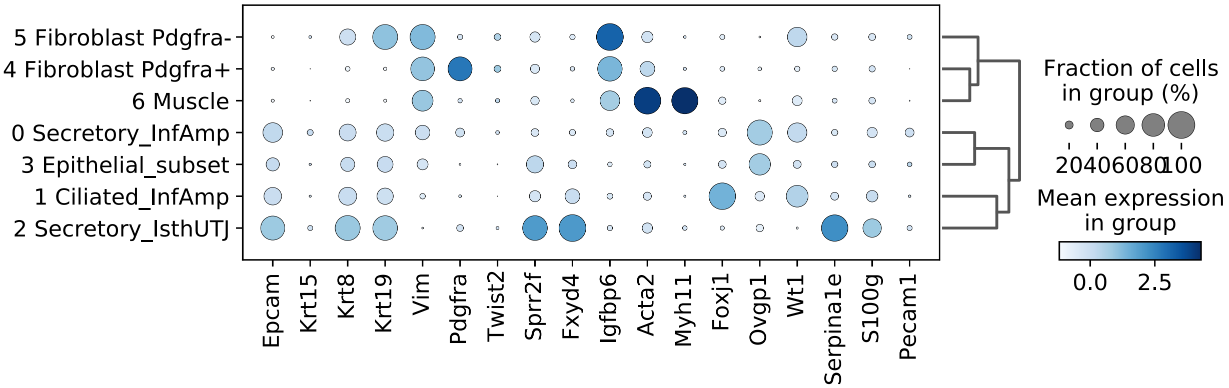


**Figure S4.** Dot plots of marker genes for each cell cluster from estrus and Veh- vs. $E_2$-treated datasets combined, including 0/secretory (InfAmp), 1/ciliated (InfAmp), 2/secretory (IsthUTJ), 3/epithelial (subset), 4/fibroblast *Pdgfra*$^+$, 5/fibroblast *Pdgfra*$^-$, and 6/muscle cell clusters.

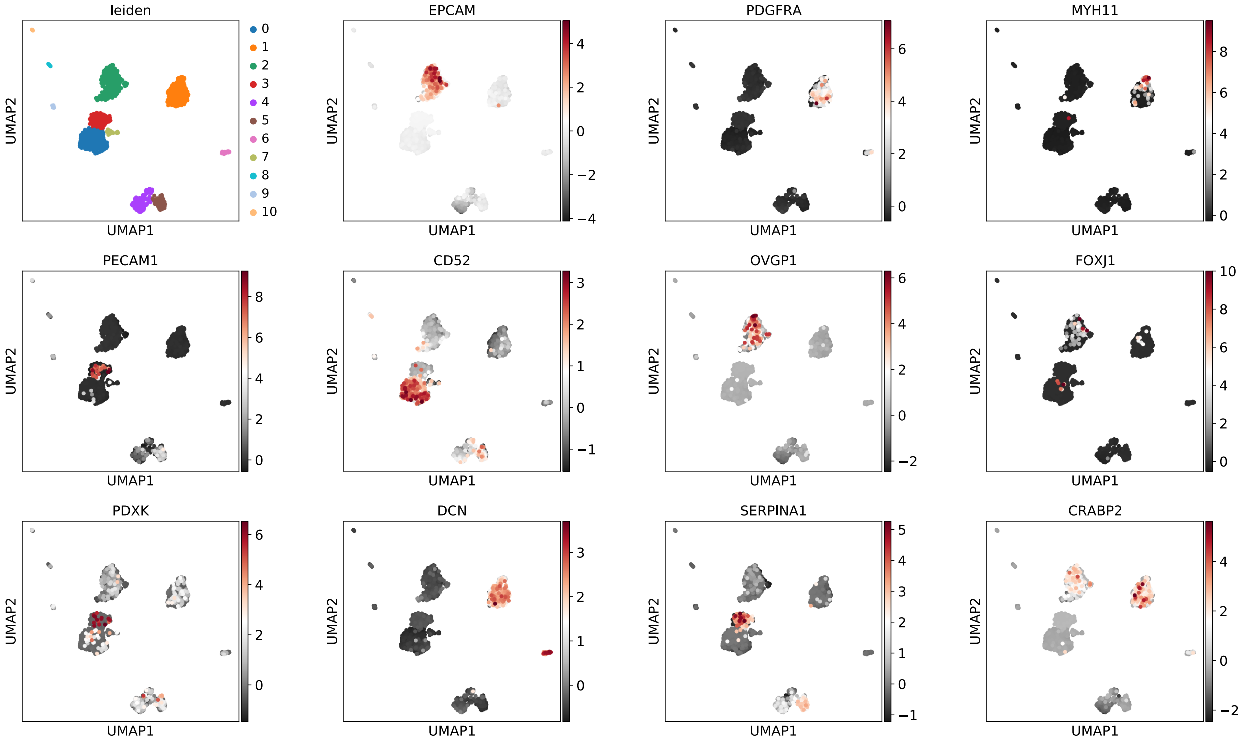


**Figure S5.** UMAP plot of clusters of cells from human fallopian tubes with different cell-type markers; epithelial cells (*EPCAM*$^+$), fibroblast (*PDGFRA*$^+$), muscle cells (*MYH11*), endothelial cells (*PECAM1*$^+$), immune cells (*CD52*$^+$), secretory epithelial cells (*OVGP1*$^+$), ciliated cells (*FOXJ1*$^+$), epithelial cells in the infundibulum and the ampulla identified in mouse oviducts (*PDXK*$^+$), fibroblast and muscle cells identified in mice (*DCN*$^+$), secretory epithelial cells in the isthmus identified in mice (*SERPINA1*$^+$), epithelial cells in the UTJ identified in mice (*CRABP2*$^+$).

### Supplementary Tables:

**Table S1:** Top 25 genes and *p*-values (*p*) enriched in each cell cluster identified in Veh- and $E_2$-treated samples; 0/ciliated (InfAmp), 1/secretory (InfAmp), 2/secretory (IsthUTJ), 3/ fibroblast *Pdgfra*$^{-}$, 4/muscle, 5/epithelial (subset), and 6/fibroblast *Pdgfra*$^{+}$ clusters.

| 0/ gene | 0/ *p* | 1/ gene | 1/ *p* | 2/ gene | 2/ *p* | 3/ gene | 3/ *p* | 4/ gene | 4/ *p* | 5/ gene | 5/ *p* | 6/ gene | 6/ *p* |
|---|---|---|---|---|---|---|---|---|---|---|---|---|---|
| *Ccdc153* | 0 | *Ovgp1* | 0 | *Fxyd3* | 0 | *Igfbp6* | 0 | *Acta2* | 6.34E-113 | *Sprr2f* | 8.94E-95 | *Dcn* | 4.96E-67 |
| *Tppp3* | 0 | *Aldoc* | 0 | *Fxyd4* | 0 | *Upk3b* | 0 | *Tagln* | 6.74E-113 | *Car2* | 3.92E-91 | *Gsn* | 2.29E-64 |
| *Elof1* | 0 | *Ier3* | 0 | *Wfdc2* | 0 | *Dcn* | 0 | *Myl9* | 7.06E-113 | *Tmem213* | 2.96E-78 | *Lgals1* | 5.03E-47 |
| *Chchd10* | 0 | *Gstm2* | 0 | *Id2* | 0 | *Rarres2* | 0 | *Tpm2* | 1.72E-110 | *Gm28940* | 1.13E-75 | *Serpinf1* | 2.34E-45 |
| *4933434E20Rik* | 0 | *Plet1* | 0 | *Slc25a48* | 0 | *Gas6* | 0 | *Tpm1* | 3.92E-110 | *4933408J17Rik* | 1.46E-75 | *Cd63* | 3.62E-45 |
| *Mt1* | 0 | *Rpl32* | 0 | *Calb1* | 0 | *Aebp1* | 0 | *Myl6* | 1.25E-109 | *A530040E14Rik* | 1.60E-75 | *Lum* | 4.50E-44 |
| *Gm19935* | 0 | *Plat* | 0 | *Grn* | 0 | *Serping1* | 0 | *Mustn1* | 9.06E-104 | *Zfp366* | 2.43E-75 | *Mirg* | 5.74E-41 |
| *Tmem212* | 0 | *Rcn1* | 0 | *Pdzk1ip1* | 0 | *Csrp2* | 0 | *Csrp1* | 1.23E-101 | *Gm47662* | 2.67E-75 | *Myrfl* | 7.38E-41 |
| *Fam183b* | 0 | *Rpl14* | 0 | *Anpep* | 0 | *Cfh* | 0 | *Flna* | 5.70E-94 | *Npas4* | 2.86E-75 | *Olfr845* | 7.52E-41 |
| *Vpreb3* | 0 | *Krt18* | 0 | *Galm* | 0 | *Sparc* | 0 | *Sparcl1* | 2.84E-93 | *Gm14412* | 3.04E-75 | *Foxn1* | 7.52E-41 |
| *Foxj1* | 0 | *Kctd14* | 0 | *Cndp2* | 1.16E-302 | *C3* | 0 | *Myh11* | 1.88E-89 | *Gm29325* | 3.10E-75 | *E030018B13Rik* | 7.53E-41 |
| *1700016K19Rik* | 0 | *Rps5* | 0 | *Rnf128* | 6.99E-296 | *Ogn* | 0 | *Mylk* | 6.44E-89 | *Gm13963* | 3.12E-75 | *Gm29010* | 7.58E-41 |
| *1110017D15Rik* | 0 | *Emb* | 0 | *Ybx1* | 1.08E-288 | *Upk1b* | 0 | *Des* | 8.24E-88 | *Col28a1* | 3.13E-75 | *Tbr1* | 7.79E-41 |
| *Dynlrb2* | 0 | *Selenom* | 0 | *Ifitm1* | 8.76E-281 | *Nbl1* | 0 | *Lgals1* | 1.01E-87 | *Gm26945* | 3.24E-75 | *Tdrd5* | 7.86E-41 |
| *Gm867* | 0 | *Rps9* | 0 | *Cldn10* | 1.79E-274 | *Nkain4* | 0 | *Vim* | 6.19E-87 | *Gm40190* | 3.24E-75 | *Ecel1* | 7.96E-41 |
| *1700007K13Rik* | 0 | *Rps27a* | 0 | *Crabp2* | 3.98E-272 | *Cldn15* | 0 | *Cald1* | 5.18E-83 | *Gm16630* | 3.38E-75 | *AI182371* | 8.03E-41 |
| *Cfap126* | 0 | *Eef1a1* | 0 | *Gsto1* | 1.01E-271 | *Rspo1* | 0 | *Crip1* | 9.12E-83 | *Gm38096* | 3.77E-75 | *A930009A15Rik* | 8.10E-41 |
| *Calm1* | 0 | *Rpl11* | 0 | *9530014B07Rik* | 1.08E-270 | *Gas1* | 0 | *Cavin3* | 3.52E-76 | *Gpr55* | 3.77E-75 | *Gm39043* | 8.24E-41 |
| *Sntn* | 0 | *Fn1* | 0 | *4833423E24Rik* | 3.64E-268 | *B2m* | 0 | *Palld* | 3.36E-73 | *Rasl10b* | 3.86E-75 | *Zfp407* | 8.48E-41 |
| *Dnah5* | 0 | *Rps15a* | 0 | *Cd9* | 1.69E-259 | *Igf1* | 0.00E+00 | *Actb* | 6.41E-73 | *Gm5160* | 4.14E-75 | *Gng10* | 9.97E-41 |
| *Tm4sf1* | 0 | *Rpl23* | 0 | *Tspan8* | 1.46E-217 | *Col3a1* | 4.65E-307 | *Mfge8* | 1.43E-69 | *A430106G13Rik* | 4.16E-75 | *Igkv13-55-1* | 1.01E-40 |
| *Dynll1* | 0 | *Slc1a3* | 3.59E-308 | *Klf5* | 1.30E-214 | *Efemp1* | 2.51E-303 | *Igfbp7* | 3.11E-67 | *Serinc4* | 4.29E-75 | *Mcmdc2* | 1.04E-40 |
| *Rsph1* | 0 | *Rps4x* | 5.41E-306 | *Pla2g4a* | 8.55E-205 | *Cavin1* | 1.55E-295 | *Malat1* | 7.06E-64 | *Slc47a1* | 4.40E-75 | *Gm40621* | 1.09E-40 |
| *Nudt4* | 0 | *Cd81* | 6.17E-305 | *Sox17* | 1.03E-202 | *Tmsb10* | 9.40E-288 | *Calm2* | 4.00E-60 | *Gm33280* | 4.60E-75 | *Gm40910* | 1.10E-40 |
| *Dnajc7* | 0 | *Rpl10* | 4.42E-303 | *Car2* | 7.82E-195 | *Crip1* | 1.26E-285 | *Actn1* | 2.39E-59 | *Rgs6* | 4.80E-75 | *Slc13a1* | 1.10E-40 |

**Table S2:** Top 500 genes and *p*-values differentially expressed in Veh- vs. $E_2$-treated samples or InfAmp vs. IsthUTJ in all 7 cell clusters combined, 0/ciliated, 1 and 2/secretory, 3 and 6/fibroblast, 4/muscle, and 5/epithelial (subset) cell clusters. Please refer to file Table S2.xlsx.

**Table S3:** Gene ontology (GO) biological processes (BPs) enriched in Veh- vs. $E_2$-treated samples or InfAmp vs. IsthUTJ in all 7 cell clusters combined, 0/ciliated, 1 and 2/secretory, and 3 and 6/fibroblast. Data were analyzed using the top 500 genes identified in Table S2. PANTHER overrepresentation test was used as analysis type with Fisher's test, FDR correction was included when applicable. There were no significant BPs enriched in 4/muscle and 5/epithelial (subset) cell clusters. Please refer to file Table S3.xlsx.

**Table S4:** Top 1,000 genes and GOBPs enriched in endogenous (estrus) vs. exogenous $E_2$-treated samples in all 7 cell clusters combined. Data were analyzed using the top 1,000 genes identified. PANTHER overrepresentation test was used as analysis type with Fisher's test and FDR correction. Please refer to file Table S4.xlsx.

**Table S5:** Top 25 genes and *p*-values enriched in 11 (#0-10) cell clusters identified from cells collected from human Fallopian tubes.

| 0/ gene | 0/ *p* | 1/ gene | 1/ *p* | 2/ gene | 2/ *p* | 3/ gene | 3/ *p* | 4/ gene | 4/ *p* | 5 gene | 5/ *p* | 6/ gene | 6/ *p* | 7/ gene | 7/ *p* | 8/ gene | 8/ *p* | 9/ gene | 9/ *p* | 10/ gene | 10/ *p* |
|---|---|---|---|---|---|---|---|---|---|---|---|---|---|---|---|---|---|---|---|---|---|
| *IL7R* | 2.27E-179 | *SPARCL1* | 2.29E-182 | *WFDC2* | 1.82E-165 | *KRTAP5-AS1* | 1.76E-130 | *HLA-DPB1* | 7.25E-81 | *LYZ* | 1.19E-63 | *GSN* | 1.37E-21 | *FTH1* | 4.46E-15 | *PTCRA* | 2.68E-13 | *CCL21* | 2.42E-11 | *IFI27* | 2.40E-07 |
| *CD69* | 1.52E-151 | *SFRP4* | 3.18E-176 | *CLU* | 7.30E-165 | *HMMR* | 3.25E-130 | *HLA-DPA1* | 2.68E-80 | *AC020656.1* | 4.38E-63 | *CFD* | 2.07E-21 | *SRGN* | 5.89E-15 | *GZMB* | 2.75E-13 | *CLDN5* | 2.64E-11 | *EGFL7* | 2.85E-07 |
| *KLRB1* | 1.01E-140 | *DCN* | 4.79E-170 | *KRT18* | 2.83E-161 | *BNIPL* | 6.17E-130 | *HLA-DQA1* | 6.77E-76 | *CSTA* | 1.62E-57 | *DCN* | 7.21E-21 | *EMP3* | 1.50E-14 | *PPP1R14B* | 2.84E-13 | *CAVIN2* | 2.92E-11 | *LIFR* | 3.15E-07 |
| *BTG1* | 3.61E-124 | *C11orf96* | 6.22E-163 | *ELF3* | 9.24E-153 | *RNF217-AS1* | 8.52E-130 | *HLA-DRB1* | 1.43E-72 | *S100A9* | 2.03E-57 | *IGFBP6* | 7.73E-21 | *SCNN1G* | 1.32E-13 | *JCHAIN* | 2.85E-13 | *TFF3* | 2.94E-11 | *CAV1* | 3.39E-07 |
| *CD52* | 2.12E-120 | *IGFBP4* | 2.72E-162 | *CLDN4* | 3.19E-147 | *TMEM255A* | 9.70E-130 | *CD74* | 6.17E-71 | *TYROBP* | 2.19E-53 | *SERPING1* | 9.13E-21 | *DLGAP2* | 1.35E-13 | *IGKC* | 2.94E-13 | *TFPI* | 4.17E-11 | *MT1M* | 3.76E-07 |
| *CD3E* | 4.62E-112 | *IGFBP7* | 1.93E-161 | *KRT8* | 1.06E-146 | *TEPP* | 1.45E-129 | *HLA-DRA* | 1.01E-70 | *CTSS* | 1.94E-52 | *MGP* | 3.92E-20 | *SLC30A2* | 1.37E-13 | *MZB1* | 3.28E-13 | *SPTBN1* | 8.16E-11 | *EMP1* | 4.09E-07 |
| *RPS12* | 3.31E-110 | *LGALS1* | 1.11E-160 | *SLPI* | 7.82E-145 | *PRC1* | 1.46E-129 | *HLA-DMA* | 3.01E-69 | *FTL* | 1.01E-50 | *FBLN5* | 6.66E-20 | *NCAPH* | 1.40E-13 | *PLD4* | 3.48E-13 | *S100A10* | 1.97E-10 | *RNASE1* | 6.45E-07 |
| *CD3D* | 5.79E-110 | *SFRP1* | 2.89E-158 | *CRISP3* | 3.56E-123 | *WNT7A* | 1.57E-129 | *HLA-DQB1* | 1.09E-67 | *TKT* | 3.00E-49 | *PLAC9* | 1.04E-19 | *S100A5* | 1.56E-13 | *IRF7* | 3.54E-13 | *KLF4* | 5.98E-10 | *NFIB* | 6.53E-07 |
| *TSC22D3* | 1.14E-109 | *CD81* | 2.95E-155 | *KRT19* | 3.66E-119 | *IL11* | 1.82E-129 | *MS4A6A* | 6.83E-60 | *LGALS2* | 3.47E-49 | *LTBP4* | 1.80E-19 | *ASPM* | 1.57E-13 | *ITM2C* | 4.44E-13 | *HLA-E* | 7.85E-10 | *SOCS3* | 7.44E-07 |
| *HARS2* | 4.66E-106 | *TPM2* | 3.92E-154 | *ELN-AS1* | 5.18E-116 | *AL137077.2* | 2.25E-129 | *GPR183* | 4.66E-59 | *S100A8* | 5.06E-49 | *CPE* | 2.98E-19 | *ZHX1* | 1.58E-13 | *CLIC3* | 5.29E-13 | *KLF2* | 1.15E-09 | *GNG11* | 7.46E-07 |
| *LINC02315* | 5.22E-106 | *RARRES2* | 4.73E-153 | *AC020909.3* | 7.50E-116 | *AC097359.2* | 2.86E-129 | *SRGN* | 7.18E-59 | *PSAP* | 6.47E-48 | *SEMA3C* | 5.79E-19 | *ZNF763* | 1.58E-13 | *IRF8* | 7.06E-13 | *AKAP12* | 1.41E-09 | *TM4SF1* | 9.85E-07 |
| *RPLP1* | 5.32E-106 | *SELENOP* | 8.09E-151 | *AC119396.2* | 1.31E-115 | *ADRA2B* | 3.03E-129 | *AIF1* | 2.08E-58 | *THBS1* | 9.85E-48 | *SFRP2* | 1.50E-18 | *AF127577.4* | 1.59E-13 | *GNA15* | 1.20E-12 | *GNG11* | 1.86E-09 | *YBX3* | 1.34E-06 |
| *AC022916.1* | 8.74E-106 | *TGM2* | 1.41E-147 | *AC011632.1* | 2.10E-115 | *KCNJ4* | 3.38E-129 | *FTL* | 5.39E-58 | *AIF1* | 2.12E-47 | *TIMP3* | 1.80E-18 | *LINC00379* | 1.64E-13 | *GRASP* | 1.39E-12 | *LMO2* | 1.87E-09 | *CCDC85B* | 2.40E-06 |
| *C10orf91* | 1.06E-105 | *PTGDS* | 1.26E-145 | *EPCAM* | 3.15E-115 | *SUOX* | 3.42E-129 | *TYROBP* | 3.59E-55 | *AP1S2* | 1.20E-46 | *MFAP5* | 2.89E-18 | *CDKL2* | 1.66E-13 | *SEC61B* | 1.66E-12 | *YBX3* | 2.22E-09 | *A2M* | 2.44E-06 |
| *EDC4* | 3.12E-105 | *SERPINF1* | 6.47E-145 | *AKAP6* | 4.00E-115 | *WNT7B* | 3.71E-129 | *SGK1* | 1.45E-54 | *TYMP* | 1.74E-46 | *CCDC80* | 2.91E-18 | *AC002091.2* | 1.69E-13 | *GPR183* | 3.66E-12 | *KANK3* | 3.12E-09 | *TSC22D1* | 3.44E-06 |
| *FABP4* | 5.54E-105 | *SELENOM* | 1.49E-139 | *AL078644.2* | 1.80E-114 | *LAMP3* | 3.78E-129 | *CTSZ* | 1.77E-54 | *VCAN* | 1.74E-46 | *ADH1B* | 7.77E-18 | *CXCL9* | 1.73E-13 | *AREG* | 7.62E-12 | *MMRN1* | 4.21E-09 | *ID1* | 3.59E-06 |
| *KCNE5* | 5.71E-105 | *TAGLN* | 1.49E-135 | *AC009831.1* | 3.41E-114 | *B4GALNT3* | 4.48E-129 | *RGS10* | 7.50E-54 | *CYBB* | 1.61E-45 | *UAP1* | 1.71E-17 | *AC097634.3* | 1.73E-13 | *NR3C1* | 9.05E-12 | *NNMT* | 4.42E-09 | *GSN* | 3.63E-06 |
| *KIFC1* | 1.03E-104 | *NBL1* | 1.85E-131 | *AC073610.3* | 3.63E-114 | *TSFM* | 5.16E-129 | *HLA-DMB* | 1.32E-53 | *S100A12* | 3.58E-44 | *ACKR3* | 2.15E-17 | *PMFBP1* | 1.77E-13 | *CD74* | 9.71E-12 | *ARL4A* | 6.76E-09 | *SPRY1* | 3.67E-06 |
| *MATN4* | 1.68E-104 | *RAMP1* | 2.67E-129 | *DNAH17* | 5.65E-114 | *DCAF4* | 5.28E-129 | *OGFRL1* | 2.92E-53 | *TNFRSF1B* | 1.11E-43 | *S100A10* | 2.85E-17 | *FAM86B2* | 1.78E-13 | *A1BG* | 1.04E-11 | *ADIRF* | 7.24E-09 | *UACA* | 3.82E-06 |
| *MPDU1* | 5.28E-104 | *IGFBP6* | 2.44E-125 | *AC027237.2* | 6.17E-114 | *AC010618.3* | 5.30E-129 | *RGS1* | 9.54E-53 | *UPP1* | 8.66E-43 | *COL6A2* | 4.52E-17 | *TMEM255B* | 1.82E-13 | *PMEPA1* | 1.72E-11 | *ANXA2* | 8.43E-09 | *SOX17* | 4.04E-06 |
| *KCNB2* | 5.77E-104 | *IFITM3* | 9.18E-125 | *SLC35F1* | 9.72E-114 | *AC051619.5* | 5.46E-129 | *MS4A7* | 1.12E-50 | *MAFB* | 1.94E-41 | *COL1A2* | 4.60E-17 | *ZBTB12* | 1.83E-13 | *PLP2* | 1.74E-11 | *IGFBP7* | 1.07E-08 | *ID3* | 6.49E-06 |
| *AP000787.1* | 8.87E-104 | *LAPTM4A* | 3.76E-124 | *IL13* | 1.34E-113 | *SPATA12* | 5.58E-129 | *FCER1G* | 2.49E-50 | *HLA-DRA* | 5.73E-41 | *IGFBP5* | 4.66E-17 | *RSPO3* | 1.88E-13 | *LDLRAD4* | 2.00E-11 | *LAYN* | 1.13E-08 | *THBD* | 7.44E-06 |
| *AC007620.2* | 9.43E-104 | *PGRMC1* | 3.63E-123 | *TACSTD2* | 1.48E-113 | *TMEM254-AS1* | 5.69E-129 | *CD83* | 6.93E-48 | *LAPTM5* | 7.01E-41 | *PMP22* | 5.47E-17 | *AC118553.1* | 1.88E-13 | *RPS3A* | 3.06E-11 | *RAB11A* | 1.13E-08 | *ADIRF* | 1.01E-05 |
